# K_2P_ channel C-type gating involves asymmetric selectivity filter order-disorder transitions

**DOI:** 10.1101/2020.03.20.000893

**Authors:** Marco Lolicato, Andrew M. Natale, Fayal Abderemane-Ali, David Crottès, Sara Capponi, Ramona Duman, Armin Wagner, John M. Rosenberg, Michael Grabe, Daniel L. Minor

**Author notes:** Equal contribution (Marco Lolicato, Andrew M. Natale). Department of Molecular Medicine, University of Pavia, Pavia ITALY (Marco Lolicato). Department of Industrial and Applied Genomics, IBM AI and Cognitive Software Organization, IBM Almaden Research Center, San Jose, CA, USA; NSF Center for Cellular Construction, University of California San Francisco, San Francisco, CA, USA.

## Abstract

K_2P_ channels regulate nervous, cardiovascular, and immune system functions^1,2^ through the action of their selectivity filter (C-type) gate^3-6^. Although structural studies show K_2P_ conformations that impact activity^7-13^, no selectivity filter conformational changes have been observed. Here, combining K_2P_2.1 (TREK-1) X-ray crystallography in different potassium concentrations, potassium anomalous scattering, molecular dynamics, and functional studies, we uncover the unprecedented, asymmetric, potassium-dependent conformational changes underlying K_2P_ C-type gating. Low potassium concentrations evoke conformational changes in selectivity filter strand 1 (SF1), selectivity filter strand 2 (SF2), and the SF2-transmembrane helix 4 loop (SF2-M4 loop) that destroy the S1 and S2 ion binding sites and are suppressed by C-type gate activator ML335. Shortening the uniquely long SF2-M4 loop to match the canonical length found in other potassium channels or disrupting the conserved Glu234 hydrogen bond network supporting this loop blunts C-type gate response to various physical and chemical stimuli. Glu234 network destabilization also compromises ion selectivity, but can be reversed by channel activation, indicating that the ion binding site loss reduces selectivity similar to other channels^14^. Together, our data establish that C-type gating occurs through potassium-dependent order-disorder transitions in the selectivity filter and adjacent loops that respond to gating cues relayed through the SF2-M4 loop. These findings underscore the potential for targeting the SF2-M4 loop for the development of new, selective K_2P_ channel modulators.

---

Selectivity filter (C-type) gating occurs in all potassium channel classes and displays a hallmark sensitivity to external potassium due to its dependency on interactions between the permeant ion and selectivity filter structure^4,6, 15-19^. Despite the fact that C-type gating is the principal K_2P_ gating mechanism^3-6^ and that previously determined K_2P_ structures show major conformational changes that affect function^7-13^, all show identical, canonical selectivity filter conformations and lack changes that could be attributed to C-type gating (Fig. S1). Notably, these structures were all determined in the presence of 150-200 mM permeant ions. In striking contrast, structure determination of a crystallizable K_2P_2.1 (TREK-1) construct, K_2P_2.1_cryst_^7^, under a series of seven potassium concentrations, 0, 1, 10, 30, 50, 100, 200 mM [K^+^] at resolutions of 3.9Å, 3.4Å, 3.5Å, 3.3Å, 3.6Å, 3.9Å, and 3.7Å, respectively, revealed obvious potassium-dependent changes in the selectivity filter structure, particularly in SF2 and the SF2-M4 loop (Figs. 1A, S2 and S3, Table S1). These changes manifested at potassium concentrations ≤50 mM and eventually encompassed all of the SF2-M4 loop and the upper portion of the selectivity filter (Gly253-Lys271) (Figs. S2 and S3). Additional changes were observed in SF1 residues Gly144-Asn147 at the lowest potassium concentrations (0 mM and 1 mM) (Fig. 1b, S3a-b). Remarkably, structure determination under the identical set of potassium concentrations in the presence of the K_2P_2.1 (TREK-1) activator ML335^7^ at resolutions of 3.4Å, 2.6Å, 3.0Å, 3.2Å, 3.2Å, 3.3Å, and 3.8Å, respectively, yielded essentially identical structures having canonical filter conformations at all potassium concentrations (Figs. 1c-d, S2, S3a-b), a result that is in line with the ability of ML335 to activate the C-type gate^7^.

**Fig. 1.**
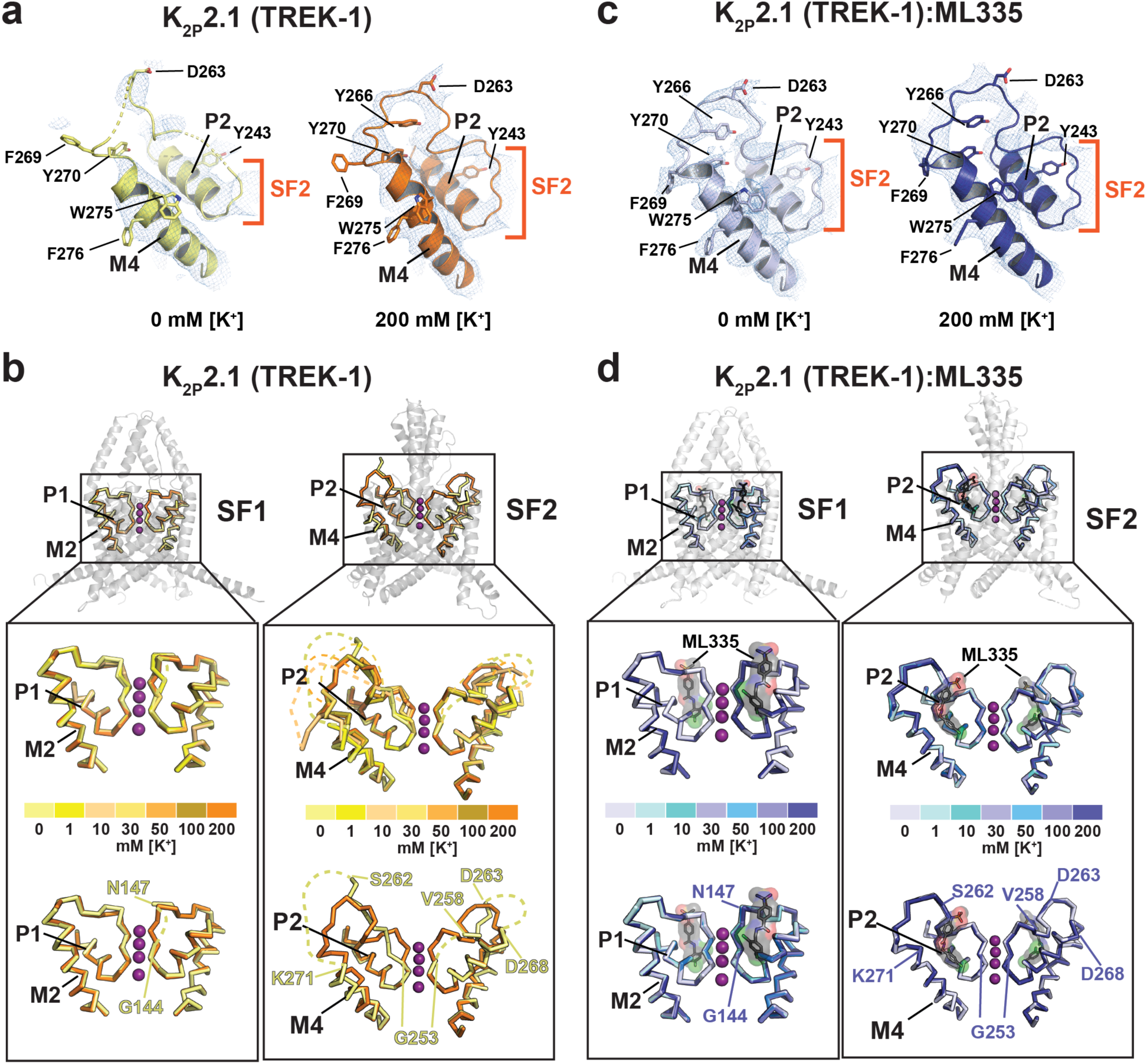
K_2P_2.1 (TREK-1) selectivity filter potassium-dependent conformational changes. **a**, Exemplar 2Fo-Fc electron density (1σ) for K_2P_2.1 (TREK-1) SF2 at 0 mM and 200 mM [K^+^]. Select residues and channel elements are indicated. Dashes indicate regions of disorder. **b**, [K^+^] dependent structural changes in K_2P_2.1 (TREK-1) SF1 (left) and SF2 (right). Top, superpositions of structures determined in 0 (pale yellow), 1 (yellow), 10 (light orange,), 30 (yellow orange), 50 (bright orange), 100 (olive), and 200 (orange) mM [K^+^]. Bottom, superposition of 0 mM and 200 mM [K^+^] structures. Dashed lines indicate regions absent from the structures. Lower panel labels mark model boundaries. **c**, Exemplar 2Fo-Fc electron density (1σ) for the K_2P_2.1 (TREK-1):ML335 complex SF2 at 0 mM and 200 mM [K^+^]. **d**, K_2P_2.1 (TREK-1):ML335 complex structural comparisons of SF1 (left) and SF2 (right). Top, superposition of structures determined in 0 (blue white), 1 (pale cyan), 10 (aquamarine), 30 (light blue), 50 (marine), 100 (slate), and 200 (deep blue) mM [K^+^]. Bottom, superposition of 0 mM and 200 mM [K^+^] structures. ML335 is black and shows its molecular surface. Lower panel labels indicate the equivalent residues from ‘**b**’. Potassium ions are from the 200 mM [K^+^] structures and are shown as magenta spheres.

Structural studies of many potassium channels have established the intimate connection between the presence of potassium ions in the selectivity filter and the conductive conformation in which the selectivity filter backbone carbonyls coordinate the permeant ions^19-23^. Hence, we asked whether the K_2P_2.1 (TREK-1) structural changes in different potassium concentrations were accompanied by changes in the number of ions in the filter. Comparison of selectivity filter region difference maps showed clear evidence for variation in the number of ions in the filter that paralleled the observed filter architecture changes. Namely, although the 100 and 200 mM [K^+^] structures showed ions at S1-S4, and the density for the S3 and S4 ions persisted even to lowest concentration examined, the densities at sites S1 and S2 were clearly absent in the 0, 1, 10, 30, and 50 mM [K^+^] structures (Figs 2a, S4a). By contrast, all of the K_2P_2.1 (TREK-1):ML335 structures showed ions at all four sites regardless of the potassium concentration, underscoring the ability of ML335 to stabilize the filter (Figs. 2b, S4b).

**Fig. 2.**
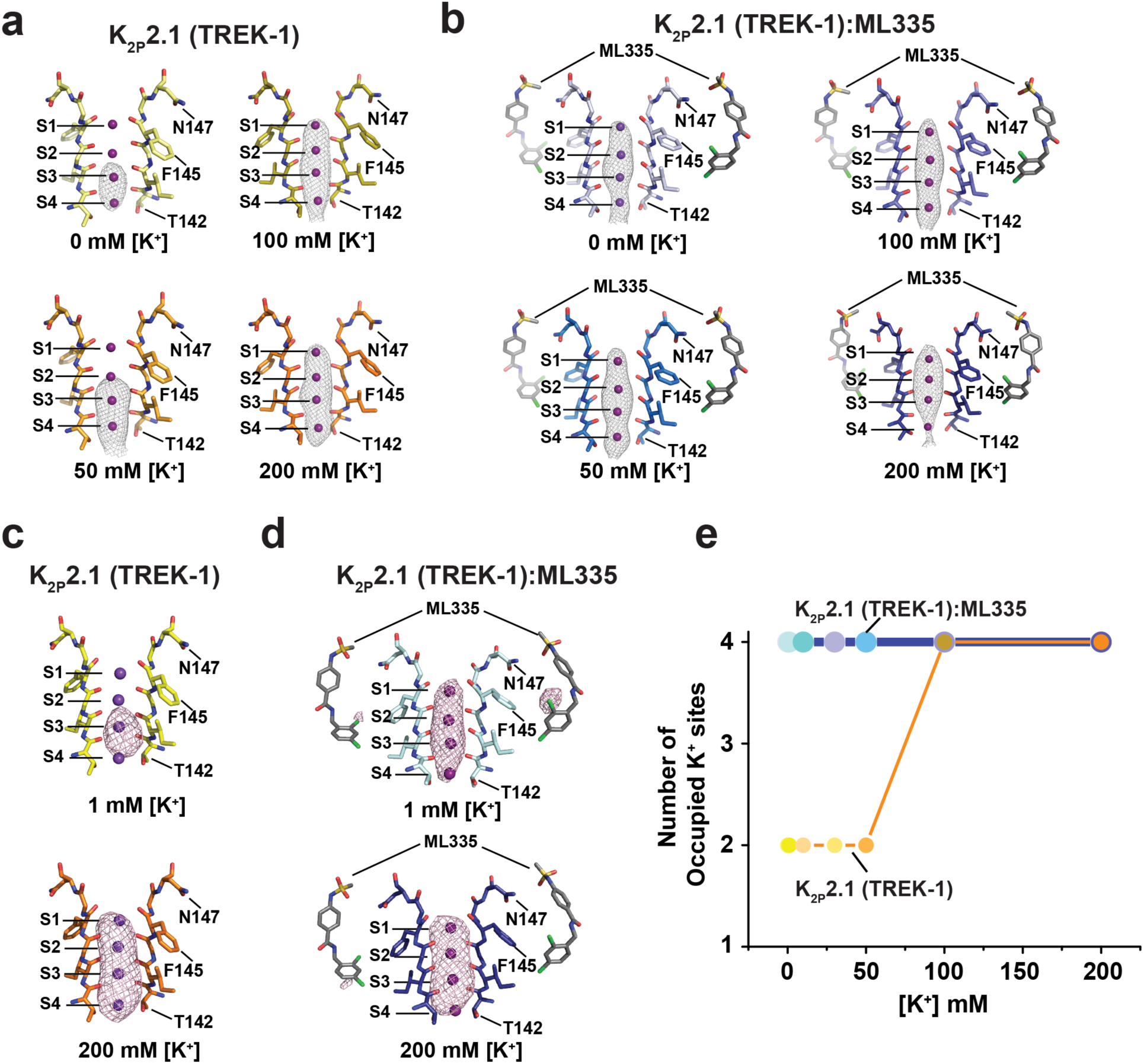
K_2P_2.1 (TREK-1) selectivity filter ion occupancy as a function of [K^+^]. **a**, and **b**, Polder omit maps^63^ for structures of **a**, K_2P_2.1(TREK-1) and **b**, K_2P_2.1(TREK-1):ML335 determined in 0 mM [K^+^] (pale yellow (5σ); blue white (4σ)), 50 mM K^+^ (bright orange (5σ); marine (4σ)), 100 mM [K^+^] (olive (4σ); slate (4σ)) and 200 mM [K^+^] (orange (4σ); deep blue (4σ)). Potassium ions are magenta spheres. Sites S1-S4 are labeled. ML335 is shown as sticks. **c**, and **d**, Potassium anomalous difference maps^24^ for **c**, K_2P_2.1(TREK-1) and **d**, K_2P_2.1(TREK-1):ML335 determined in 1 mM [K^+^] (yellow (4σ); pale cyan (4σ)) and 200 mM [K^+^] (orange (4σ); deep blue (4σ)). In ‘**a-d**’ SF1 in the 200 mM [K^+^] conformation is shown for reference. S1-S4 sites and select amino acids are labeled. **e**, Plot of the number of observed selectivity filter ions as a function of [K^+^]. Colors correspond to the scheme in Fig. 1c and d.

To confirm that the changes in the number and positions of selectivity filter ions resulted from potassium ions, we used long wavelength X-rays above and below the potassium K-absorption edge (λ=3.3509 Å and 3.4730 Å) to measure potassium anomalous scattering ^24,25^ from crystals having 1 mM [K^+^] or 200 mM [K^+^] in the absence or presence of ML335. Anomalous difference maps showed unequivocally that potassium ions occupy sites S1-S4 under 200 mM [K^+^] conditions regardless of the presence of ML335 (Fig. 2c,d). By contrast, the density from 1 mM [K^+^] conditions showed an unmistakable ML335-dependent difference in the number of potassium ions that agreed with our other observations (Fig. 2a,b, Table S2). In the absence of the activator, potassium ions were observed only in the lower portion of the filter, whereas potassium ions are found at all four positions in presence of ML335 (Fig. 2c,d). Together, these data demonstrate that the dramatic loss of structure observed in the upper portion of SF2 as potassium concentrations are lowered is indeed accompanied by a loss of potassium ions at sites S1 and S2 (Fig. 2e). These potassium-dependent structural changes are entirely suppressed by the C-type gate activator ML335. Hence, when taken with the prior demonstration that ML335 directly activates the C-type gate^7^, the conformational differences between the low [K^+^] structures with and without ML335 reflect the high and low activity states of the C-type gate, respectively.

To gain further insight into how K^+^ occupancy and ML335 impact the C-type gate, we turned to molecular dynamics (MD) simulations of K_2P_2.1 (TREK-1) using conditions of 180 mM [K^+^] and a +40 mV applied membrane potential alone (denoted ‘High [K^+^]/+40 mV’, 36.5 µs aggregate) and with ML335 (denoted ‘High [K^+^]/+40 mV/ML335’, 31.6 µs aggregate). Both conditions showed many permeation events (144 and 253 for High [K^+^]/+40 mV and ‘High [K^+^]/+40 mV/ML335’, respectively), confirming that the initial structures represent conduction competent states. Nevertheless, the pattern of permeation events with respect to time showed striking differences depending on the presence of ML335 (Fig. 3a). Over the course of the simulations, the majority (7/12) of the High [K^+^]/+40 mV trajectories entered long lived (>1 µs) non-conducting states from which they did not recover. By contrast, only 2 of 10 High [K^+^]/+40 mV/ML335 trajectories entered into long-lived, non-conducting states. Concordantly, the two conditions had a substantial difference in the median conductance (7 pS vs 32 pS for High [K^+^]/+40 mV and High [K^+^]/+40 mV/ML335, respectively) (Fig. 3b).

**Fig. 3.**
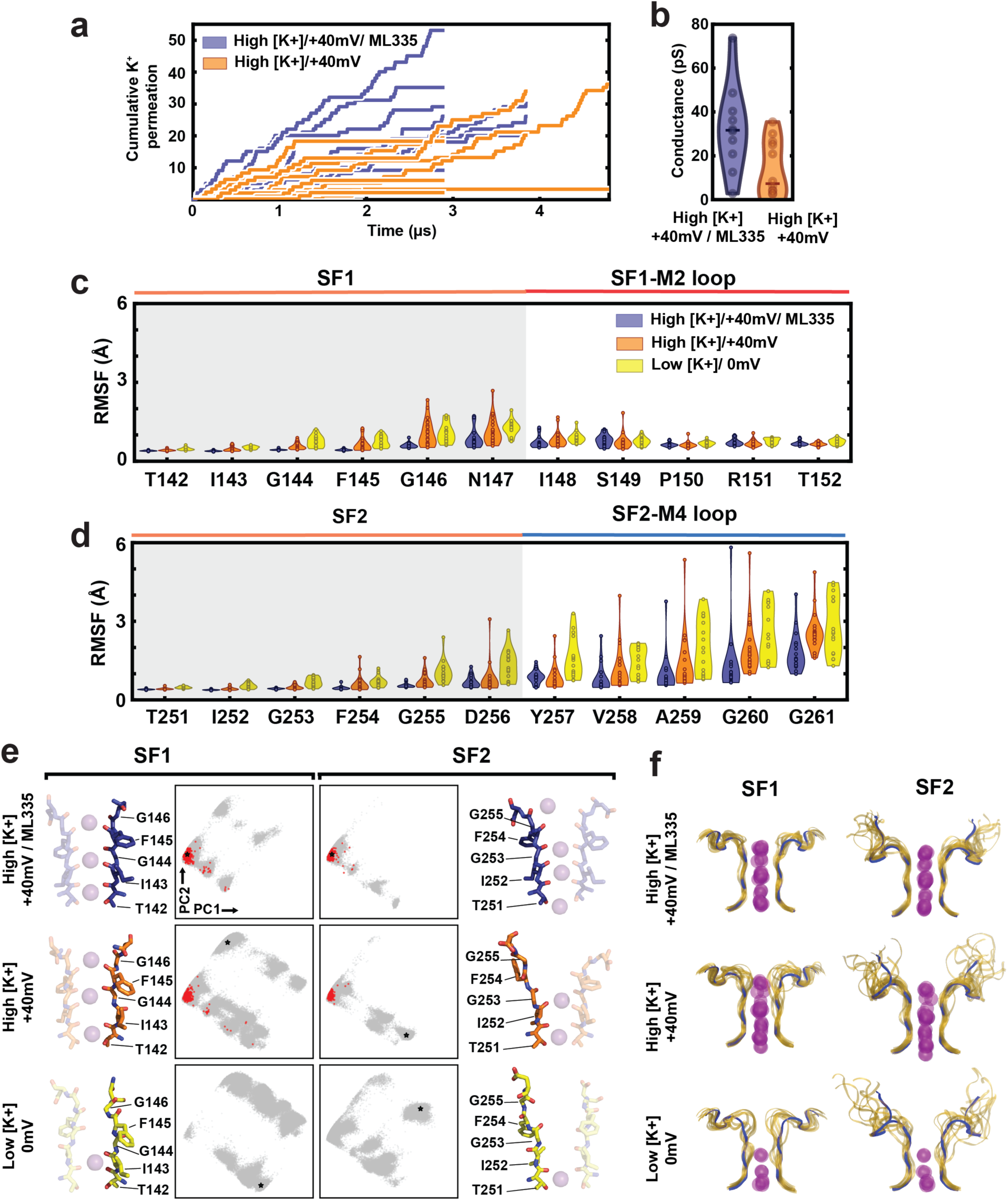
K_2P_2.1 (TREK-1) conductance properties and SF conformational dynamics from MD simulations. **a**, Cumulative K^+^ ion permeation events over simulation time for all individual trajectories in ‘High [K^+^]/+40 mV’ (orange) and ‘High [K^+^]/+40 mV/ML335’ (purple) conditions. **b**, Conductance values calculated from the trajectories in ‘**a’**. Each point shows the conductance calculated from an independent trajectory; median values for each condition are given as horizontal bars. **c**, and **d**, C_α_ root-mean-squared-fluctuation (RMSF) values of the filter and loop regions for **c**, pore domain 1, and **d**, pore domain 2, for all simulated conditions. Each point represents an RMSF value calculated from one K_2P_2.1 (TREK-1) subunit of one trajectory. Conserved selectivity filter signature sequences are shaded grey. **e**, PCA analysis of SF1 and SF2 dihedral angles and exemplar filter conformations. Each dot represents the instantaneous conformation of the TIGFG backbone dihedral angles from single selectivity filter. Black stars indicate the location in PC1-2 space of the adjacent exemplar. Red dots indicate conformations immediately preceding K^+^ permeation events. **f**, Final SF1 and SF2 and loop ion and backbone conformations from all simulation trajectories. Final frame of each individual trajectory is shown as transparent gold ribbons; potassium ions are shown as transparent purple spheres. Solid blue ribbons represent the initial crystal structure conformation.

To determine if there were differences in C-type gate dynamics across simulation conditions, we calculated root-mean-squared-fluctuation (RMSF) values for the selectivity filter and the post-filter loops. Because low potassium occupancy in the filter resulted in increased mobility in these regions of the crystal structures, we included a third set of simulations in which K_2P_2.1 (TREK-1) had only a single ion in the filter under no applied membrane potential (denoted, ‘Low [K^+^]/0 mV’, 20.6 µs aggregate). This analysis revealed that residues Phe145-Ser149 of SF1, Phe254-Gly261 of SF2, and the SF2-M4 loop comprise the three most dynamic areas near the filter and showed that their mobility was greatly restricted by ML335 (Fig. 3c-d). Further, under Low [K^+^]/0 mV conditions the mobility of these regions exceed either of the High [K^+^]/+40 mV conditions. Together, the simulations indicate that the absence of K^+^ in the filter versus the presence of ML335 have strong, opposite effects on the dynamics of the selectivity filter and SF2-M4 loop (Fig. 3c-d).

To compare selectivity filter conformations across all simulated conditions, we performed principal component analysis (PCA) on the backbone dihedral angles of the SF1 and SF2 ion-coordinating ‘TIGFG’ amino acid motifs. Projecting onto the first two principal components (PC1 and PC2) (Fig. 3e) uncovered a distinct grouping of SF1 and SF2 conformations that lack major deviations from the initial structure. All prior K_2P_ structure selectivity filters (Fig. S5a-b) (denoted as the ‘native state’) as well as selectivity filters from other potassium channels thought to capture either conducting states^21,26^ or, surprisingly, inactivated states^21,27^, map to the center of this group (Fig. S5c). Additionally, performing clustering analysis on the PC space revealed many distinct clusters of non-native conformations in which the backbone dihedral angles deviate substantially from the native state (Fig. S5d). Some of these conformations are reached from the canonical SF1 structure via a single discrete amide ‘crankshaft’ motion between either the S2/S3 sites (Ile143/Gly144) (Fig. S5d, cluster 2) or at the top of the S0 site (Gly146/Asn147) (Fig. S5d, cluster 3) and correspond to conformations suggested to be involved in C-type gating in previous K_2P_ channel simulations^28,29^. The remaining SF1 and SF2 clusters represent larger deviations from the initial model and cannot be described by single amide crankshaft motions. Some of these configurations are reminiscent of the unusual selectivity filter structure of the nonselective channel NaK^30^ (Fig. S5c-d), while others represent novel configurations that have not been observed experimentally (Fig. S5d). Notably, non-canonical conformations are more highly populated in the absence of ML335 under low [K^+^] conditions (Fig. 3e-f) and are much more dramatic than those reported for simulations of selectivity filters from four-fold symmetric potassium channels^31,32^ or K_2P_s^28,29^.

Of all of the K_2P_2.1 (TREK-1) conformations observed under high [K^+^] conditions, only a few are compatible with K^+^ permeation. Greater than 90% of ion conduction events occurred when all four SF strands occupied the native state conformational cluster. No conduction events were observed when more than two SF strands adopted non-native states. Under High [K^+^]/+40 mV/ML335 conditions, SF1 and SF2 were found in the native state cluster 90% and 95% percent of the time, respectively, while under High [K^+^]/+40 mV conditions these values dropped to 64% and 86% (Fig. 3e). Thus, the presence of ML335 reduces the accessible conformational space of the filter, restricting SF1 and SF2 largely to their native conductive conformations. This conformational restriction causes longer periods of sustained conduction and higher conductance values relative to the High [K^+^]/+40 mV condition (Fig. 3a-b), in line with the fact that ML335 directly activates the K_2P_2.1 (TREK-1) C-type gate^7^. The Low [K^+^]/0 mV simulations experienced even greater selectivity filter conformational diversity that included conformations similar to the non-conductive states observed in high [K^+^] conditions as well as more extreme non-native conformations.

In all three simulation conditions, most of the non-conductive filter conformations have multiple deviations from the canonical structure (Fig. 3f and S5d) and share a loss of ion binding sites at either S1, S2, or both due to rearrangement of the ion coordinating carbonyls (Fig. 3d bottom and middle right panels, and Fig. S5d). These changes leave only the S3 and S4 sites competent for potassium binding, and agree with the crystallographic ion positions under low potassium conditions (Figs.1-2, 3f and S2-S3). Furthermore, examination of the ensemble of final SF1 and SF2 backbone conformations from the simulations under different conditions shows that these structural components display increased conformational disorder and pseudo-four-fold symmetry breaking that is in excellent agreement with the X-ray structures (Fig. 3f). SF1 adopts non-native conformations, particularly around Asn147, that pinch the conduction pathway, whereas SF2 preferentially dilates out of the pathway (Supplementary videos 1-2). This asymmetry extends beyond the filter. The SF1-M2 loop remains largely native-like, despite the changes in SF1 while the longer SF2-M4 loop is dynamic and adopts many different conformations as SF2 changes (Supplementary video 2). This later observation explains the loss of density for this region in the low [K^+^] crystal structures (Fig. 1a,b). Taken together, the structures and simulations support the idea that ML335 acts by stabilizing the K_2P_ selectivity filter in a conductive state and indicate that the low [K^+^]crystal structures represent a C-type inactivated state in which asymmetric disorder in the extracellular portion of the selectivity filter disrupts the S1 and S2 ion binding sites and inhibits ion conduction.

In most potassium channels, including the first K_2P_ pore domain (PD1), a six-residue linker connects the extracellular end of the selectivity filter to the pore domain outer transmembrane helix (Fig. 4a-b and S6a-b). K_2P_s are unique in that the second pore domain linker (PD2) is longer than this canonical length by 6-8 residues in 14 of the 15 K_2P_ subtypes (Fig. S6c-d). Despite these differences, the N-terminal portions of the PD1 and PD2 linkers adopt very similar structures up to Pro150 and Ala259, respectively (Fig. 4a). The remaining residues form a loop supported by a conserved hydrogen bond network, the Glu234 network, comprising the Glu234 carboxylate from M3, the SF2-M4 loop Gly260 backbone amide, and the Tyr270 phenolic -OH from M4 (Fig. 4c-d).

**Fig. 4.**
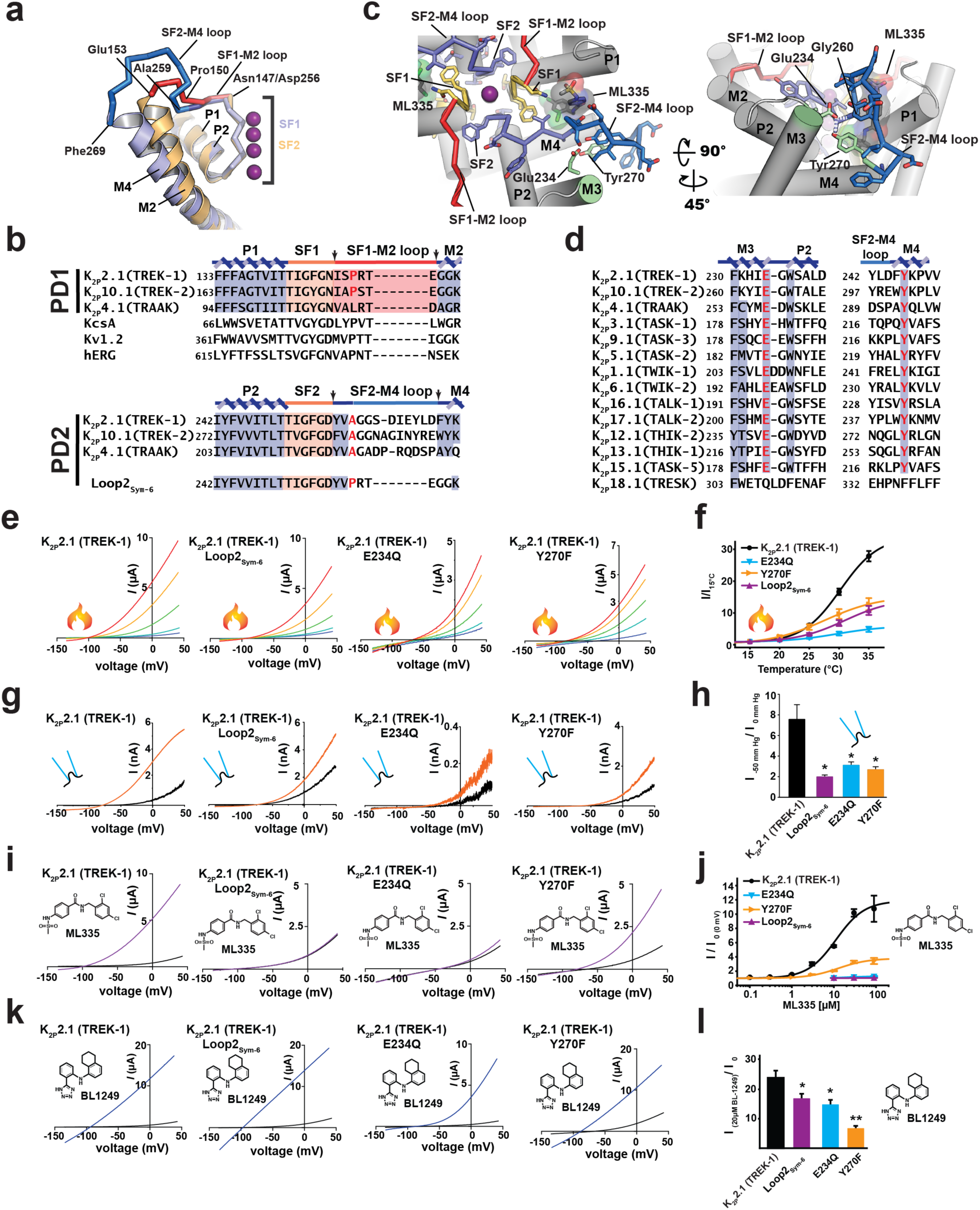
The K_2P_ SF2-M4 loop is central to C-type gate function. **a**, Superposition of the P1-SF1-M2 (orange) and P2-SF2-M4 (slate) regions of the K_2P_2.1 (TREK-1) pore domain. SF1-M2 loop (red) and SF2-M4 loop (blue) are indicated. Dark blue indicates the section of the SF2-M4 loop that is similar to the SF1-M2 loop. Residue labels indicate the beginning and end of the SF1-M2 and SF2-M4 Loops as well as the point of structural divergence (Pro150/Ala259). **b**, Sequence comparison of the SF regions of Pore domain 1 (PD1) and Pore domain 2 (PD2) for the indicated channels and select homotetrameric potassium channels. The arrows denote the ends of the selectivity filter-outer transmembrane helix linker. Pro150/Ala259 equivalents are highlighted red. **c**, Structural details of the SF2-M4 loop in the K_2P_2.1 (TREK-1):ML335 complex. SF1 (yellow), SF2 (slate), SF1-M2 (red), and SF2-M4 (marine) are indicated. Conserved hydrogen bond network of Glu234 (green), Tyr270 (green), Gly260 (marine) is indicted. ML335 is shown in both stick and space filling rendering. **d**, Conservation of the Glu234-Tyr270 pair among human K_2P_ channels: K_2P_2.1 (TREK-1) AAD47569.1, K_2P_10.1 (TREK-2) NP_612190.1, K_2P_4.1 (TRAAK) AAI10328.1, K_2P_3.1 (TASK-1) NP_002237.1, K_2P_9.1 (TASK-3) NP_001269463.1, K_2P_5.1 (TASK-2) NP_003731.1, K_2P_1.1 (TWIK-1) NP_002236.1, K_2P_6.1 (TWIK-2) NP_004814.1, K_2P_16.1 (TALK-1) NP_115491.1, K_2P_17.1 (TALK-2) AAK28551.1, K_2P_12.1 (THIK-2) NP_071338.1, K_2P_13.1 (THIK-1) NP_071337.2, K_2P_15.1 (TASK-5) EAW75900.1, K_2P_18.1 (TRESK) NP_862823.1. SF1 and SF2 sequence and numbers for K_2P_2.1 (TREK-1)_cryst_ (PDB:6CQ6)^7^ are identical to that of K_2P_2.1 (TREK-1) AAD47569.1. Glu234 and Tyr270 positions are highlighted red. **e**, Exemplar two-electrode voltage clamp recordings of the responses of the indicated channels at 15°C (blue), 20°C (light green), 25°C (lime green), 30°C (orange, and 35°C (red). **f**, Normalized temperature responses (n≥10). **g**, Exemplar inside-out recordings of the responses of the indicated channels to pressure. 0 mm Hg (black). 50 mm Hg (orange). **h**, Averaged pressure responses for the indicated channels (n≥4). **i**, Exemplar two-electrode voltage clamp recordings of the responses of the indicated channels to 30 µM ML335 (purple). **j**, ML335 dose-response curves for the indicated channels (n≥3). EC_50_ 11.3 ± 3.4 and 12.7 ± 4.1 µM, Max activation 11.9 ± 1.3 and 3.8 ± 0.4 fold for K_2P_2.1 (TREK1) and K_2P_2.1 (TREK1) Y270F, respectively. **k**, Exemplar two-electrode voltage clamp recordings of the responses of the indicated channels to 20 µM BL-1249 (blue). **l**, Normalized responses to 20 µM BL-1249 for the indicated channels (n≥7). In panels **f, h, j**, and **l**, channels are indicated as K_2P_2.1 (TREK1) (black), K_2P_2.1 (TREK1) Loop2_sym6_ (purple), K_2P_2.1 (TREK1) E234Q (light blue), and K_2P_2.1 (TREK1) Y270F (orange). ‘*’ and ‘**’ indicate p < 0.05 and p < 0.001, respectively.

When bound to the K_2P_ modulator pocket^7^, ML335 interacts with the side of the SF2-M4 loop opposite to the Glu234 network (Fig. 4c) and strongly attenuates potassium-dependent loop dynamics (Figs. 1b-d, 3c-d, S2, and S3). Further, in the simulations both potassium and ML335 greatly influence the Glu234 network integrity (Figs. S6e-f). Therefore, we set out to test consequences of restricting the SF2-M4 loop mobility and disrupting the Glu234 network.

To create a channel having symmetric length loops between each selectivity filter and its outer transmembrane helix, we transplanted Pro150-Gly155 from PD1 onto PD2, denoted ‘Loop2_Sym-6_’ (Fig. 4b). Loop2_Sym-6_ showed blunted responses to temperature (Fig.4 e,f) and pressure (Fig. 4 g,h). Consistent with the deletion of key ML335 binding SF2-M4 loop residues, Loop2_Sym-6_ was unresponsive to ML335 (Fig. 4 i,j), but remained partially sensitive to BL-1249, an activator that affects the channel from a site under the selectivity filter^33,34^. Importantly, measurement of rectification in inside-out patches, a parameter that is a direct measure of C-type gate activation^5,7^, demonstrated that unlike gain-of-function mutants^7^, Loop2_Sym-6_ does not have a constitutively activated C-type gate that would render it insensitive to gating commands (Fig. S7a-b). Hence, the blunted responses caused by shortening the SF2-M4 loop to the canonical length indicate that the unusual length of the SF2-M4 loop is central to C-type gate control.

Disruption of the hydrogen bond network by E234Q and Y270F mutations resulted in channels having severely blunted responses to temperature (Fig. 4e-f), pressure (Fig. 4g-h), ML335 (Fig. 4i-j), and BL-1249 (Fig. 4k-l). Unlike Loop2_Sym-6_, both mutations compromised ion selectivity as evidenced by an altered reversal potential (Figs. 3e, g, i, k, and S8). This baseline selectivity defect was partially corrected by temperature or pressure activation (Fig. 3g, j, and S8). Importantly, inside-out patch clamp experiments demonstrated that neither mutant resulted in channels having a C-type gate that was activated at rest, even though Y270F caused a slight decrease of the rectification coefficient (Fig. S7a-b). Unexpectedly, we also found that E234Q exhibited a time and voltage dependent inactivation (Fig. S7c-d), further validating the importance of the Glu234 network for C-type gate control. Together, these data strongly support the key role that the SF2-M4 loop has in K_2P_ channel gating.

Despite the central role of the selectivity filter C-type gate in K_2P_ channel function^3-6^, observation of conformational changes therein has eluded previous structural studies^7-13,35^. Our data establish that control of the K2P C-type gate involves unprecedented, asymmetric, potassium dependent order-disorder transitions in the selectivity filter and surrounding loops. This conformational change eliminates the S1 and S2 ion binding sites (Fig. 5, Supplementary videos 3-4) and consists of two classes of selectivity filter changes. One exposes the SF1 Asn147 sidechains to the extracellular solution and pinches the selectivity filter extracellular side (Fig. 5a, Supplementary video 3). The second unwinds SF2 and the SF2-M4 loop and widens the selectivity filter (Fig. 5b, Supplementary video 4). Such asymmetric changes could contribute to the bimodal distribution of closed state dwell times reported for K_2P_2.1 (TREK-1)^36^ and the closely-related K_2P_10.1 (TREK-2)^37^. The structural rearrangements and loss of S1 and S2 ions, together with the demonstration that destabilization of the SF2-M4 loop structure compromises ion selectivity are reminiscent of the changes that convert the nonselective bacterial channel NaK having only the S3 and S4 sites into a potassium-selective channel capable of binding ions at sites S1-S4^14^ and agree with the loss of ion selectivity associated with K_2P_ C-type gating^4,18,38^.

**Fig. 5.**
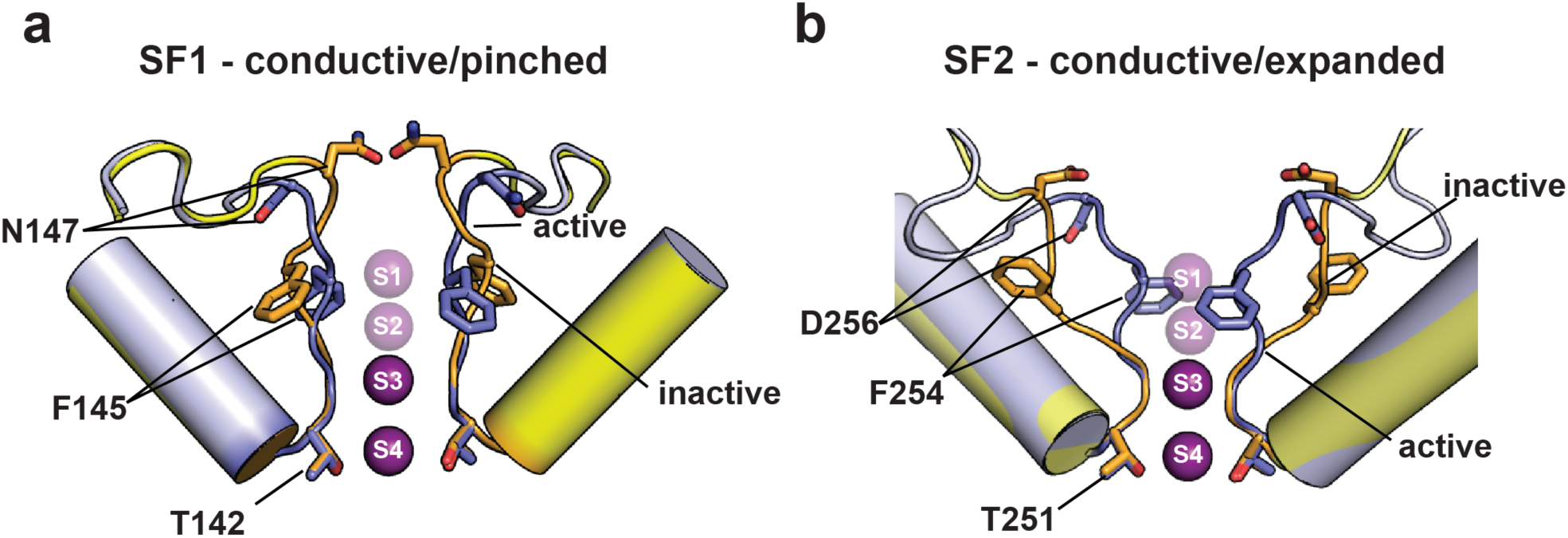
Structural changes associated with C-type gating. **a**, SF1 (yellow) and **b**, SF2 (light blue) selectivity filter changes between the active (**a**, and **b**, conductive) and inactive (**a**, pinched and **b**, expanded) conformations based on the 1 mM [K^+^] and 0 mM [K^+^]:ML335 structures, respectively. Selectivity filters for 1 mM [K^+^] (yellow orange) and 0 mM [K^+^]:ML335 (slate) show select residues. Potassium ions are magenta spheres.

The SF1 structural transformation affects a post-filter residue Asn147, a position that modulates C-type inactivation in four-fold symmetric potassium channels^39,40^ and that undergoes similar changes in hERG simulations^41^. Hence, this class of C-type inactivation conformational change appears to be shared with four-fold symmetric potassium channels. The substantial differences in the degree of conformational changes between SF1 and SF2 appear to depend on the loop length connecting these elements to the outer transmembrane helix of their respective pore domains and indicate that the dramatic SF2 conformational changes result from the uniquely long K_2P_ SF2-M4 loop. Binding of small molecules to the K_2P_ modulator pocket enables conduction by stabilizing the SF2-M4 loop and selectivity filter, whereas, disruption of the integrity of the SF2-M4 loop blunts transduction of gating cues that originate from the intracellular C-terminal tail^3,42-47^ and pass through M4 to the C-type gate^3,4^. These findings support the ideas that the K_2P_ selectivity filter and its supporting architecture are dynamic under basal conditions^7^, that ion permeation requires restrictions of filter conformations^7^, that permeant ions organize and stabilize the K_2P_ conductive state^5,34^, and that the position of M4 is not the sole K_2P_ activation determinant^7,48^. The demonstration that the SF2-M4 loop has a key role in transducing gating cues sensed by other channel components to the K_2P_ selectivity filter gate highlights the potential for targeting this unique K_2P_ loop for selective small molecule or biologic modulators directed at K_2P_-dependent processes such as anesthetic responses^49,50^, pain^51-53^, arrythmia^54^, ischemia^49,55,56^, and migraine^57^.

## Methods

### Protein expression and purification

An engineered mouse K_2P_2.1 (TREK-1), denoted K_2P_2.1_cryst_, encompassing residues 21–322 and bearing the following mutations: K84R, Q85E, T86K, I88L, A89R, Q90A, A92P, N95S, S96D, T97Q, N119A, S300A, E306A, a C-terminal green fluorescent protein (GFP), and His_10_ tag was expressed and purified from *P. pastoris* as previously described^7^.

### Crystallization and refinement

Purified K_2P_2.1_cryst_ was concentrated to 6 mg ml^-1^ by centrifugation (Amicon Ultra-15, 50 kDa molecular mass cut-off; Millipore) and crystallized by hanging-drop vapor diffusion at 4 °C using a mixture of 0.2 μl of protein and 0.1 μl of precipitant over 100 μl of reservoir containing 20–25% PEG400, 200 mM KCl, 1 mM CdCl_2_, 100 mM HEPES, pH 8.0. Crystals appeared in 12 h and grew to full size (200–300 μM) in about one week.

Crystals were harvested and cryoprotected with buffer D (200 mM KCl, 0.2% OGNG, 15 mM HTG, 0.02% CHS, 1 mM CdCl_2_, 100 mM HEPES, pH 8.0,) with 5% step increases of PEG400 up to a final concentration of 38%. After cryoprotection, crystals were incubated for 8 hours in buffer E (38% PEG400, 0.2% OGNG, 15 mM HTG, 0.02% CHS, 1 mM CdCl_2_, 100 mM HEPES, pH 8.0,) to which has been added a combination of NaCl and KCl up to 200mM salt to achieve the following K^+^ concentrations: 0 mM, 1 mM, 10 mM, 30 mM, 50 mM, 100 mM and 200 mM. In the soaking experiments where the activator was present, ML335 was added to the soaking cocktail to a 1 mM final concentration. Crystals were subsequently harvested and flash-frozen in liquid nitrogen.

Datasets for K_2P_2.1_cryst_ in the presence of differing potassium concentrations, alone or with ML335, were collected at 100 K using synchrotron radiation at APS GM/CAT beamline 23-IDB/D Chicago, Illinois, processed with XDS^58^, scaled and merged with Aimless^59^. Final resolution cut-offs were 3.9Å, 3.5Å, 3.4Å, 3.3Å, 3.6Å, 3.9Å and 3.7Å for K2P2.1cryst in the presence of 0 mM, 1 mM, 10 mM, 30 mM, 50 mM, 100 mM and 200 mM potassium, respectively, using the CC_1/2_ criterion. Final resolution cut-offs for the K_2P_2.1_cryst_:ML335 complex were 3.4Å, 2.6Å, 3.0Å, 3.2Å, 3.2Å, 3.3Å and 3.8Å in the presence of 0 mM, 1 mM, 10 mM, 30 mM, 50 mM, 100 mM and 200 mM potassium, respectively. Structures were solved by molecular replacement using the K_2P_2.1_cryst_ structure (PDB: 6CQ6)^7^ as search model purged of all the ligands. Several cycles of manual rebuilding, using COOT^60^, and refinement using REFMAC5^61^ and PHENIX^62^ were carried out to improve the electron density map. Two-fold local automatic NCS restraints were employed during refinement. Two potassium ions were modeled into 2Fo-Fc densities of the Apo K_2P_2.1_cryst_ 0 mM, 1 mM, 10 mM, and 50 mM structures; whereas, four potassium ions were modeled into 2Fo-Fc densities of the Apo K_2P_2.1_cryst_ 100 mM and 200 mM structures. Four potassium ions were modeled for all the K_2P_2.1_cryst_:ML335 complexes. To validate the presence of the potassium ions, a polder map^63^ was generated for each structure. The polder map of the Apo K_2P_2.1_cryst_ 50 mM structure showed a density in the filter that extended beyond the S3 site into the S2 site; however, modeling an additional low occupancy K^+^ ion at this site did not improve the overall statistics. Attempts to refine the occupancy of this third ion using PHENIX^62^ yielded an ion having zero occupancy. Hence, the final structure has two ions in the filter, even though there may be a low occupancy ion present that is not accountable due to the resolution limit of the data. The final cycle of refinement of each structure was carried out using BUSTER^64^.

### K^+^ anomalous data collection

Long-wavelength data were collected at beamline I23, Diamond Light Source ^25^, UK at a temperature ∼50 K at wavelengths of 3.3509 Å and 3.4730 Å, above and below the potassium K absorption edge, processed and scaled with XDS/XSCALE^58^. Anomalous difference Fourier maps to locate the potassium positions were calculated with ANODE^65^ using the K_2P_2.1 (TREK-1) structure (PDB:6CQ6)^7^. Peaks present in the maps above, but absent in the maps below the absorption edge were assigned as potassium.

### Two-electrode voltage-clamp electrophysiology

Two-electrode voltage-clamp recordings were performed on defolliculated stage V–VI *Xenopus laevis* oocytes 18–48 h after microinjection with 1-40 ng cRNA. Oocytes were impaled with borosilicate recording microelectrodes (0.3–3.0 MΩ resistance) backfilled with 3 M KCl. Recording solution (96 mM NaCl, 2 mM KCl, 1.8 mM CaCl_2_, and 2.0 mM MgCl_2_, buffered with 5 mM HEPES, pH 7.4, except where otherwise indicated) was perfused by gravity.

Currents were evoked from a -80 mV holding potential followed by a 300 ms ramp from -150 mV to +50 mV. Data were acquired using a GeneClamp 500B amplifier (MDS Analytical Technologies) controlled by pClamp software (Molecular Devices), and digitized at 1 kHz using Digidata 1332A digitizer (MDS Analytical Technologies).

For temperature experiments, recording solutions were heated by an SC-20 in-line heater/cooler combined with an LCS-1 liquid cooling system operated by the CL-100 bi-polar temperature controller (Warner Instruments). Temperature was monitored using a CL-100-controlled thermistor placed in the bath solution 1 mm upstream of the oocyte. Perfusate was warmed from 15°C to 35°C in 5°C increments, with recordings performed once temperature readings stabilized at the desired values. Temperature response data were fit with the equation 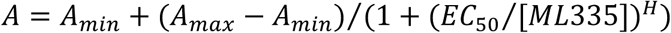 where A_min_ and A_max_ are the minimum and maximum activation, respectively, T_1/2_ is the temperature of half maximal activation, and S is the slope factor^4^.

Dose response experiments were conducted at room temperature (22°C) and used standard recording solution supplemented with 0.2% DMSO and the indicated concentration of ML335^7^. Dose response data were fit with the equation *A* = *A*_*min*_ + (*A*_*max*_ − *A*_*min*_)/(1 + (*EC*_50_/[*ML*335])^*H*^) where A_min_ and A_max_ are the minimum and maximum activation, respectively, EC_50_ is the half maximal effective concentration, and H is the Hill slope^7^. Data analysis and curve fitting were performed using Clampfit and python according to procedures adapted from^4,7^.

### Patch clamp electrophysiology

Human embryonic kidney cells (HEK293) were grown at 37°C under 5% CO_2_ in a Dulbecco’s modified Eagle’s medium (DMEM) supplemented with 10% fetal bovine serum (FBS), 10% L-glutamine, and antibiotics (100 IU Ml^−1^ penicillin and 100 mg mL^-1^ streptomycin). Cells were transfected (in 35 mm diameter wells) using Lipofectamine 2000 (Invitrogen) and a pIRES-GFP (Invitrogen) plasmid vector into which the gene encoding for mouse K_2P_2.1 (TREK-1) wild type or mutants has been inserted in the first cassette ^4^. 1 µg DNA was used for K_2P_2.1 (TREK-1) and Loop2-sym-6, whereas 3 µg DNA was necessary to record reliable currents from E234Q and Y270F. Transfected cells were identified visually using the green fluorescent protein (GFP) expressed in the second cassette of the pIRES-GFP plasmid vector. After a minimum of 6 hours post-transfection, cells were plated onto coverslips coated with Matrigel (BD Biosciences).

The inside-out configuration of the patch clamp technique was used to record K^+^ or Rb^+^ currents at room temperature (23 ± 2°C) 24–48 h post-transfection^5,7^. Data acquisition was performed using pCLAMP 10 (Molecular Devices) and an Axopatch 200B amplifier (Molecular Devices). Pipettes were pulled from borosilicate glass capillaries (TW150F-3; World Precision instruments) and polished (MF-900 microforge; Narishige) to obtain 1–2 MΩ resistances.

Stretch activation of K_2P_2.1 (TREK-1) and mutants was performed by applying a -50 mmHg pressure to the inside-out patch through a High-Speed Pressure Clamp (HSPC-1, ALA Scientific Instruments) connected to the electrode suction port, after recording the current at 0 mm Hg. Pipette solution contained 150 mM NaCl, 5 mM KCl, 1 mM CaCl_2_, 2 mM MgCl_2_, and 20 mM HEPES (pH 7.4 with NaOH). Bath solution contained 145 mM KCl, 3 mM MgCl_2_, 5 mM EGTA, and 20 mM HEPES (pH 7.2 with KOH) and was continuously perfused at 200 ml hr^-^1 during the experiment. K_2P_2.1 (TREK-1) currents were elicited by a 1 s ramp from -140 to +50 mV from a - 80 mV holding potential.

Voltage-dependent activation and inactivation of K_2P_2.1 (TREK-1) and mutants were recorded from inside-out patches. Pipette solution contained 150 mM KCl, 3.6 mM CaCl_2_, and 10 mM HEPES (pH 7.4 with KOH). Bath solution contained 150 mM RbCl, 2 mM EGTA, and 10 mM HEPES (pH 7.4 with RbOH), and was continuously perfused at 200 ml hr^-1^ during the experiment. For voltage-dependent activation, currents were elicited by voltage steps from -100 mV to +100 mV, from a –80 mV holding potential. For voltage-dependent inactivation, currents were elicited by pre-pulse voltage steps from -50 mV to +90 mV from a –80 mV holding potential, each step being followed by a test pulse at +100 mV. All electrophysiology data were analyzed using Clampfit 9 (Molecular Devices).

### Molecular Dynamics

#### Simulation setup

Initial K_2P_2.1 (TREK-1) simulations in the absence of ML335 were initiated from PDB:5VK5. Later simulations were based on PDB:6CQ6^7^, which is indistinguishable from PDB:5VK5 except for a minor difference in the C-terminal portion of M4. Simulations in complex with ML335 were constructed from PDB:6W8C. In both cases, models consisted of residues 35-321, a disulfide bond was formed between C93 in one subunit with C93 in the other, missing loops were built with RosettaRemodel^66^, and N- and C-termini were capped with methylamide and acetyl groups, respectively. Structures were embedded in pure 1-palmitoyl-2-oleoyl-glycero-3-phosphocholine (POPC) or POPC + 1-stearoyl-2-arachidonoyl-glycero-3-phosphatidylinositol 4,5-bisphosphate (PIP_2_) containing bilayers using CHARMM-GUI ^67^ and solvated in 180 mM [K^+^] with neutralizing Cl^-^ (excepting low [K^+^] simulations, which contained only 4 mM [K^+^]). The K^+^ ion placement in the selectivity filter under high K^+^ conditions was S1/S2/S4 with an empty S3 site. Low K^+^ simulations were initiated with a single ion in the selectivity filter placed at various sites (Table S3). The protein, lipids, water and ML335 were parameterized with CHARMM36m^68^, CHARMM36^69^, TIP3P^70^, and CGenFF 3.0.1^71-73^, respectively.

#### Simulation details

Production data was collected on two platforms: Anton2^74^ at the Pittsburgh Supercomputing Center and local GPU resources using GROMACS 2018^75^ (see Table S3 for a full list). All systems were equilibrated locally using a similar equilibration protocol in which the protein and ligand heavy atoms were initially restrained with a 5 kcal/mol/Å^2^ force constant that was gradually reduced over 10-12 ns. Next, for systems simulated under a membrane potential, we performed a 10 mV voltage jump every 5 ns until reaching 40 mV using the constant electric field protocol^76^. Note that systems destined for Anton2 were equilibrated with NAMD 2.13^77^. Production run details varied by hardware. Simulations on Anton2 used a 2.5 fs timestep, an MTK barostat^78^ with semi-isotropic pressure control at 1 atm, and a Nose-Hoover thermostat^79,80^ with a temperature of 303.15 K. Additionally, nonbonded interactions were cut off at 10 Å, long-range electrostatic interactions were calculated using the Gaussian split Ewald method^81^, and hydrogens were constrained with SHAKE ^82^. Meanwhile, GROMACS 2018 runs used either a 2 or 2.5 fs timestep, a Parrinello-Rahman barostat^83,84^ with semi-isotropic pressure control at 1 atm, and a Nose-Hoover thermostat set to 310 K. Nonbonded interactions were cut off at 12 Å with force-switching between 10-12 Å, long-range electrostatics were calculated with particle mesh Ewald^85^, and hydrogens constrained with LINCS algorithm^86^. For low K^+^ simulations, solution ions were excluded from the selectivity filter using a flat bottom restraint on Anton2 or harmonic positional restraints in GROMACS 2018.

#### Simulation analysis

Ions were tracked within a 22 Å long cylindrical volume centered on the selectivity filter, and a permeation event was recorded when an ion originating below (above) the mid-plane of the filter (defined by the plane separating S2-S3 sites) exited the top (bottom) of the cylinder. The time of the permeation event was recorded as the last time the ion crossed the mid-plane prior to exit from the cylinder. Principal component analysis was carried out on the backbone dihedral angles of selectivity filter residues (142-146 in SF1 and 251-255 in SF2) as described in^87^ and each strand was treated independently. Formation of hydrogen bonds to carboxylate or carbonyl oxygens was determined on the basis of the H to O distance (>2.5 Å for OH donors, >2.75 Å for NH donors) and C=O H angle >110°. For all analyses, conformations were sampled from the trajectories every 480-500 ps. All analysis code was built on top of the MDAnalysis python package^88^.

## Supporting information

Supplementary Video 1 High [K+]/+40 mV simulation trajectory.

Supplementary Video 2 Low [K+]/0 mV simulation trajectory.

Supplementary Video 3 Morph between the SF1 active and inactive conformations.

Supplementary Video 4 Morph between the SF2 active and inactive conformations.

## Acknowledgements

We thank C. Arrigoni and L. Pope for electrophysiology guidance, C. Colleran for cell culture assistance, L. Jan for support, and K. Brejc for comments on the manuscript. This work was supported by grants NIH-R21NS091941 to J.M.R., R01GM089740 to M.G., and NIH-R01-MH093603 to D.L.M., and an AHA postdoctoral fellowship to F.A.-A..

## Author Contributions

M.L., A.M.N., F.A.-A., S.C., J.M.R., M.G., and D.L.M. conceived the study and designed the experiments. M.L. expressed, purified, and crystallized the proteins, collected diffraction data, and determined the structures. R.D and A.W. collected anomalous diffraction data. M.L., A.M.N., F.A.-A., and, D.C. performed functional studies. A.M.N., S.C., J.M.R., and M.G. designed and executed the simulations. M.L., A.M.N. F.A.-A., D.C., M.G., and D.L.M. analyzed the data. M.G. and D.L.M analyzed data and provided guidance and support. M.L., A.M.N., F.A.-A., M.G. and D.L.M. wrote the paper.

## Competing interests

The authors declare no competing interests.

## Materials and correspondence

Correspondence should be directed to M.G. or D.L.M. Requests for materials should be directed to D.L.M.

## Data and materials availability

Coordinates and structures factors are deposited in the RCSB and will be released immediately upon publication.

## Supplementary Materials

**Fig. S1.**
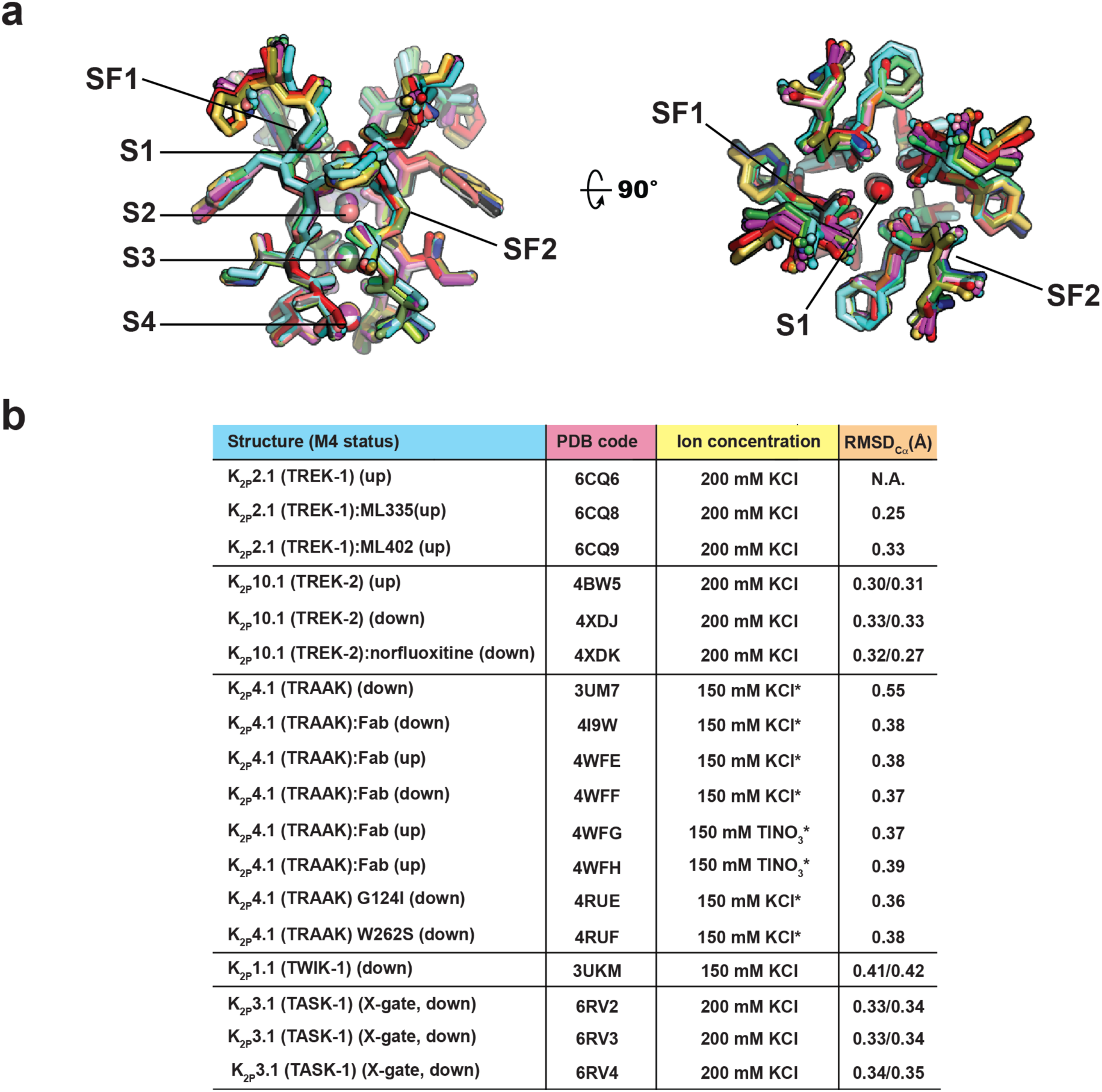
K_2P_ channel selectivity filters structure comparison. **a**, Superposition of the selectivity filters and permeant ions for: K_2P_2.1 (TREK-1) (6CQ6)^7^ (smudge), K_2P_2.1 (TREK-1):ML335 (6CQ8)^7^ (deep salmon), K_2P_2.1 (TREK-1):ML402 (6CQ9)^7^; K_2P_10.1 (TREK-2) (4BW5)^8^ (pink), (4XDJ)^8^ (magenta), (4XDK)^8^ (purple); K_2P_4.1 (TRAAK) (3UM7)^9^ (aquamarine), (4I9W)^10^ (limon), (4WFE) (forest green)^11^, (4WFF) (white)^11^,(4WFG) (grey)^11^, (4WFH) (black)^11^; K_2P_4.1 (TRAAK) G124I (4RUE) (blue)^12^ K_2P_4.1 (TRAAK) W262S (4RUF) (lime green)^12^; K_2P_1.1 (TWIK-1) (3UKM)^13^ (red). K_2P_3.1 (TASK-1) (6RV2) (orange)^35^, K_2P_3.1 (TASK-1):BAY1000493 (6RV3) (yellow orange)^35^, and K_2P_3.1 (TASK-1):BAY2341237(6RV4) (olive)^35^. SF1, SF2 and ion binding positions, S1-S4, are indicated. Ions are shown as spheres and colored according to the parent structure. **b**, K_2P_ channel structures, permeant ion concentration in crystallization conditions, and RMSD for all selectivity filter backbone atoms relative to K_2P_2.1 (TREK-1) (6CQ6)^7^. Structures with two RMSD values indicate structures having chains A/B and C/D, respectively. ‘*’ indicates samples where permeant ions were part of the protein sample buffer.

**Fig. S2.**
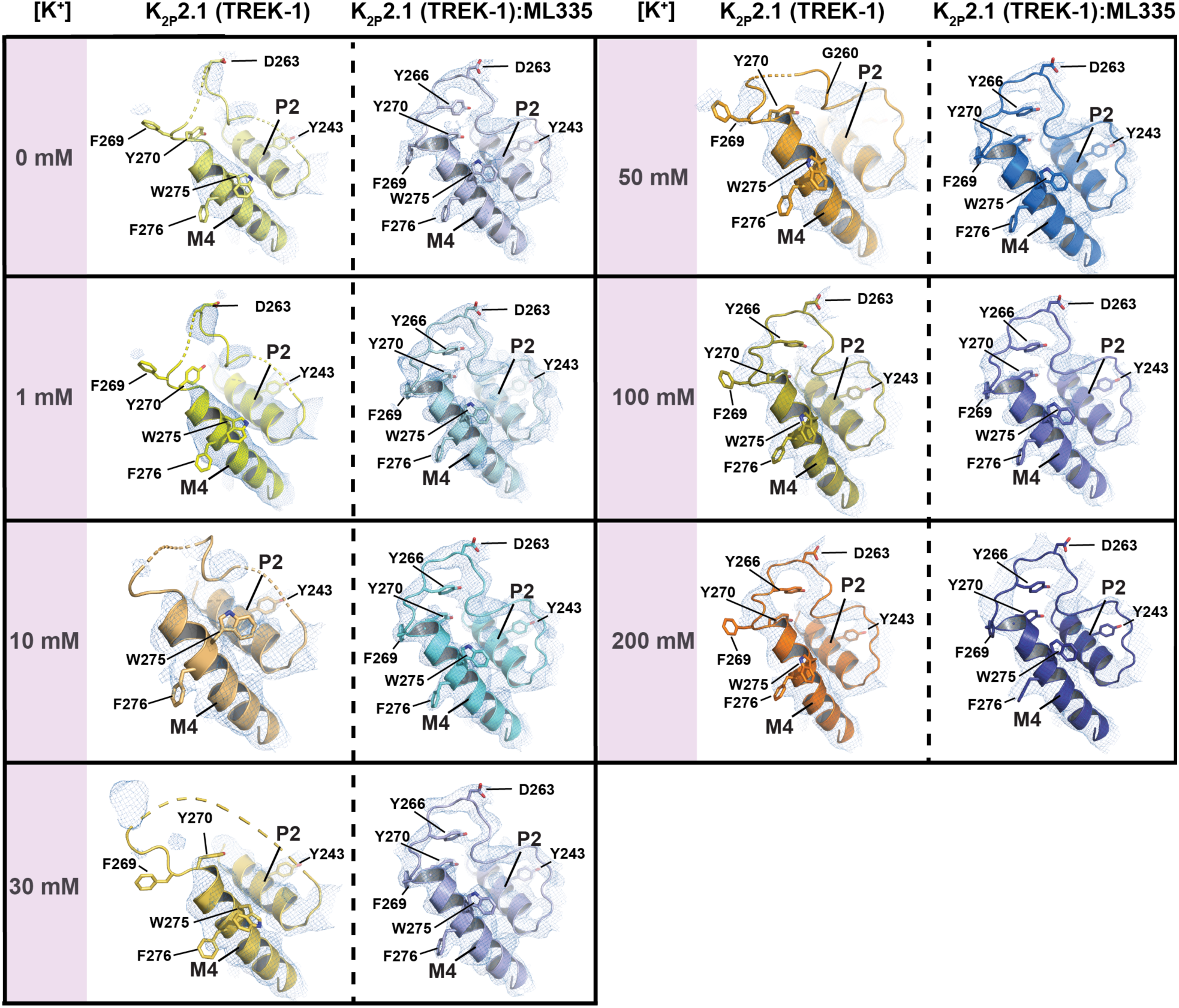
K_2P_2.1 (TREK-1) selectivity filter potassium-dependent conformational changes. SF2 exemplar 2Fo-Fc electron density (1σ) for K_2P_2.1 (TREK-1) (left) and K_2P_2.1 (TREK-1):ML335 (right) structures under 0 mM (pale yellow; blue white), 1 mM (yellow; pale cyan), 10 mM (light orange; aquamarine), 30 mM (yellow orange; light blue), 50 mM (bright orange; marine), 100 mM (olive; slate), and 200 mM (orange; deep blue) [K^+^]. Dashed lines indicate regions of disorder. Select residues and channel elements are indicated.

**Fig. S3.**
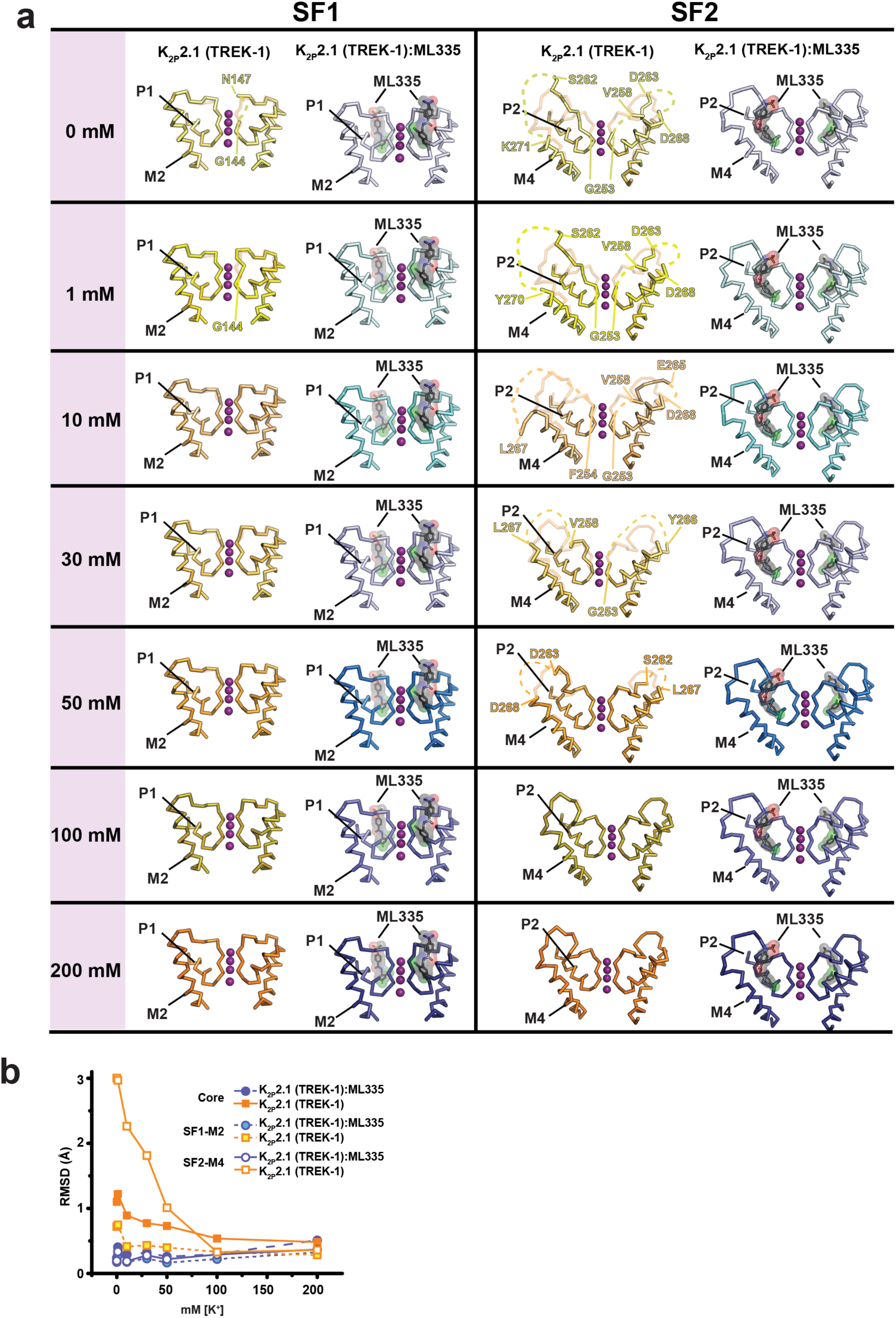
Selectivity filter structural changes as a function of potassium concentration. **a**, K_2P_2.1 (TREK-1) and the K_2P_2.1 (TREK-1):ML335 complex SF1 and SF2 structures determined at the indicated potassium concentrations: 0 mM (pale yellow; blue white), 1 mM (yellow; pale cyan), 10 mM (light orange; aquamarine), 30 mM (yellow orange; light blue), 50 mM (bright orange; marine), 100 mM (olive; slate), and 200 mM [K^+^] (orange; deep blue). K_2P_2.1 (TREK-1) panels show an overlay with the 200 mM [K^+^] K_2P_2.1 (TREK-1) structure in lighter shading. Labels indicate the last visible residue at points where the chain becomes disordered. Potassium ions from the 200 mM [K^+^] structures are shown in all panels as a reference. **b**, K_2P_2.1 (TREK-1) and K_2P_2.1 (TREK-1):ML335 complex RMSD_Cα_ as a function of [K^+^]. Structures are compared to K_2P_2.1 (TREK-1) in 200 mM [K^+^] (PDB:6CQ6) and K_2P_2.1 (TREK-1):ML335 complex in 200 mM [K^+^] (PDB:6CQ8), respectively. Channel elements are grouped as follows: Core: residues 50-146, 153-255, 269-311; SF1-M2: residues 142-188; and SF2-M4: residues 251-295.

**Fig. S4.**
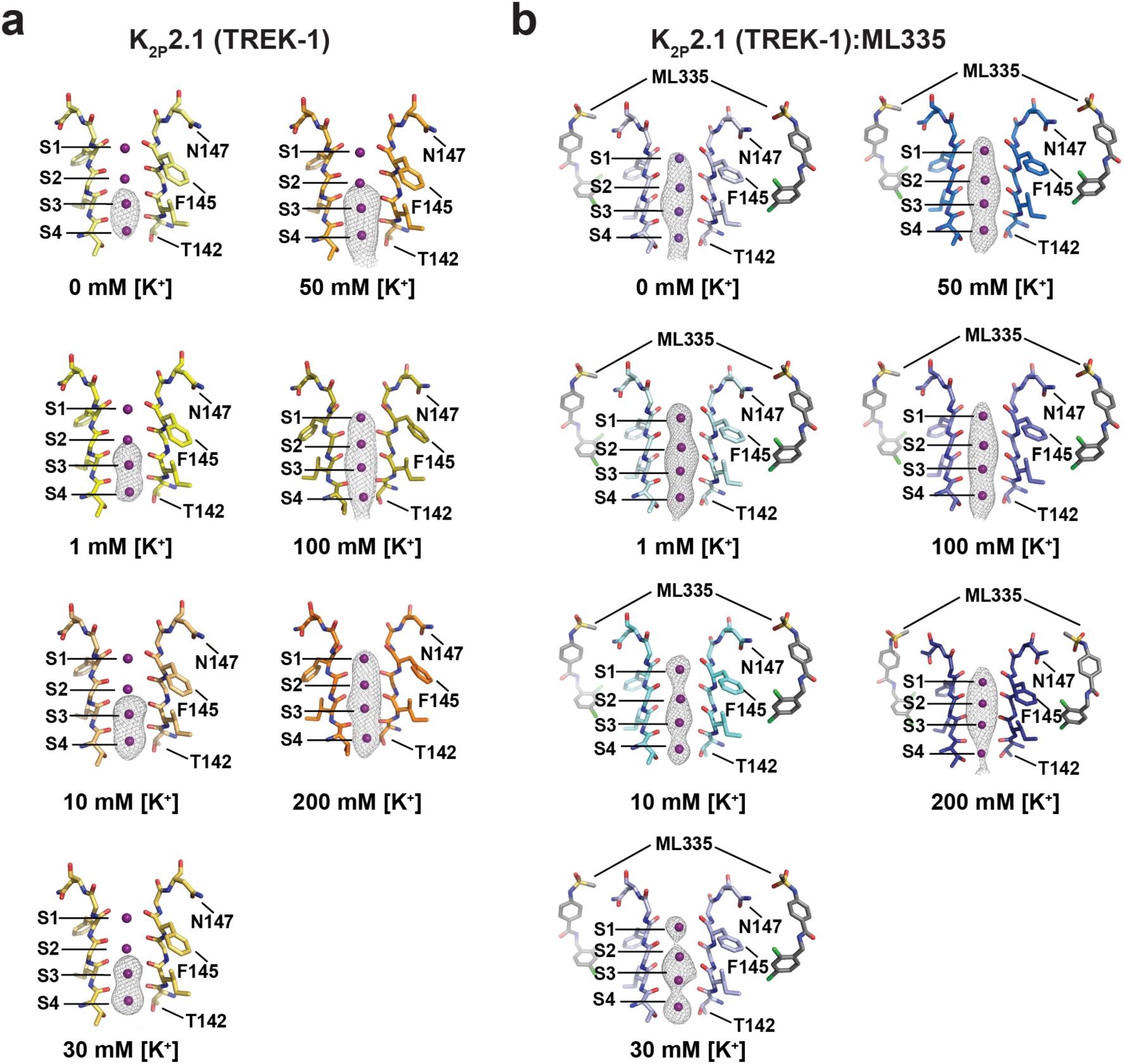
Omit maps showing K2P2.1 (TREK-1) selectivity filter ion occupancy as a function of [K^+^]. **a,b**, Polder omit maps^63^ for structures of K_2P_2.1 (TREK-1) determined in 0 mM [K^+^] (pale yellow) (5σ), 1 mM [K^+^] (yellow) (4σ), 10 mM [K^+^] (light orange) (5σ), 30 mM [K^+^] (yellow orange) (4σ), 50 mM [K^+^] (bright orange)(5σ), 100 mM [K^+^] (olive) (4σ), and 200 mM [K^+^] (orange)(4σ) (**a**) or K_2P_2.1 (TREK-1):ML335 determined in 0 mM [K^+^] (blue white) (4σ), 1 mM [K^+^] (pale cyan) (4σ), 10 mM [K^+^] (aquamarine) (4σ), 30 mM [K^+^] (light blue) (4σ), 50 mM [K^+^] (marine) (4σ), 100 mM [K^+^] (slate) (4σ) and 200 mM [K^+^] (deep blue) (4σ) (**b**). Potassium ions are magenta spheres. Sites S1-S4 are labeled. ML335 is shown as sticks. SF1 in the 200 mM [K^+^] conformation is shown for all panels. Select residues are indicated.

**Fig. S5.**
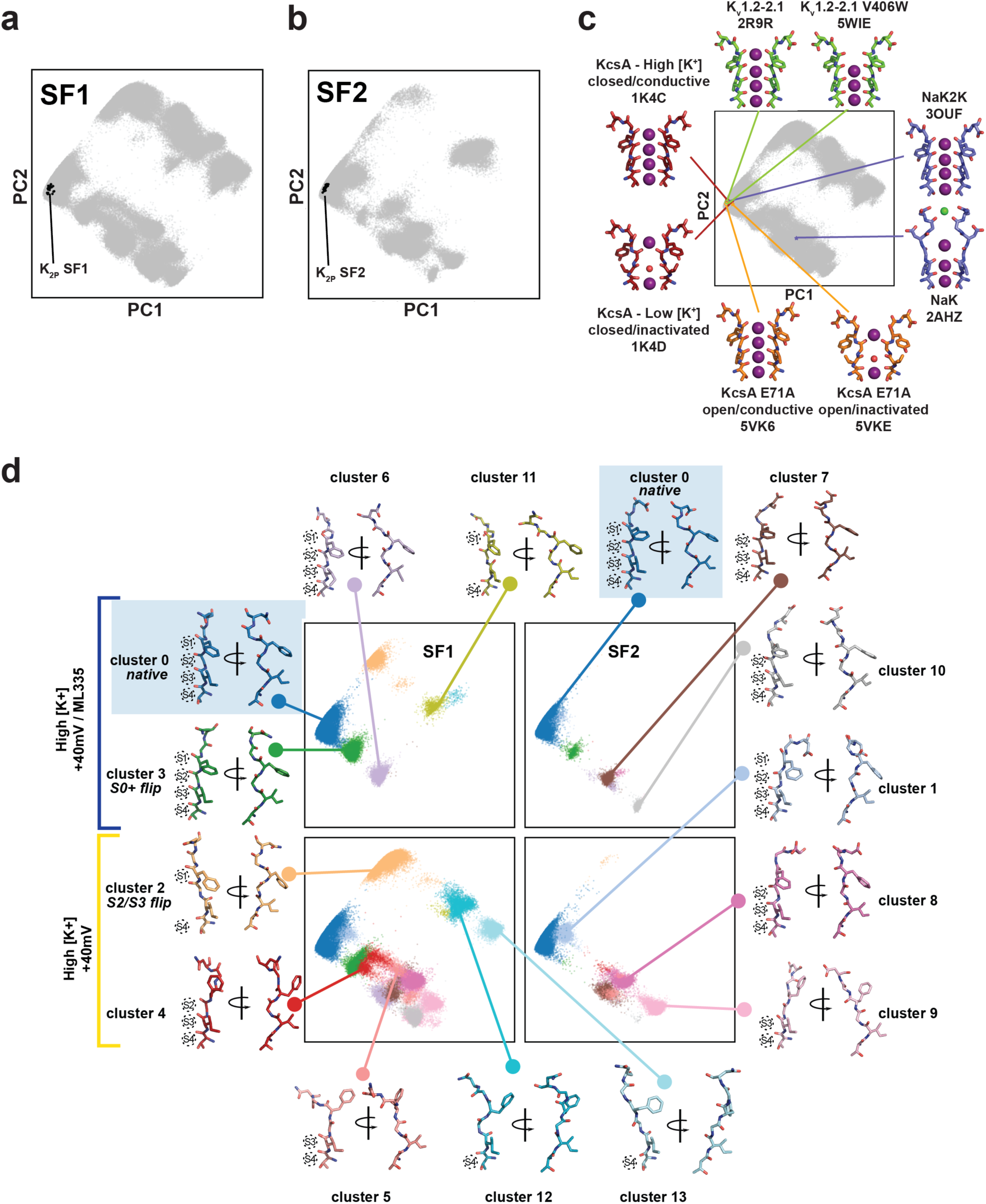
K_2P_2.1 (TREK-1) principal component analysis (PCA). **a,b**, PCA projection of selectivity filter conformations for SF1 (**a**) and SF2 (**b**). Grey points indicate K_2P_2.1 (TREK-1) conformations from all simulations in this work; black points represent projections of SF1 and SF2 conformations obtained from the Fig. S1b K_2P_ crystal structures. **c**, Representative non-K_2P_ selectivity filter conformations and their projections into PCA space. Grey points indicate K_2P_2.1 (TREK-1) conformations from all simulations (SF1 and SF2). Lines and colored points show the PCA projected location of SF conformations from the indicated tetrameric potassium channel crystal structures: KcsA closed/conductive (1K4C) and closed/inactivated (1K4D)^21^ (firebrick); Kv1.2-2.1 chimera (2R9R)^26^ and V406W mutant (5WIE)^27^ (chartreuse); NaK (2AHZ)^30^ and K^+^ selective mutant NaK2K (3OUF)^14^ (slate); and KcsA E71A open/conductive (5VK6) and open/inactivated (5VKE)^22^ (orange). **d**, Hierarchical clustering of SF conformations from all High [K^+^] simulations. Clustering was performed on PC1-3, for clarity only PC1 and PC2 are shown. Points representing selectivity filter conformations in PCA space are colored according to their membership in one of 14 identified clusters. A single representative conformation is shown for each cluster, with the exception of the native state (cluster 0) for which representative conformations from both SF1 and SF2 are shown. For each representative conformation intact ion binding sites are indicated with dotted circles.

**Fig. S6.**
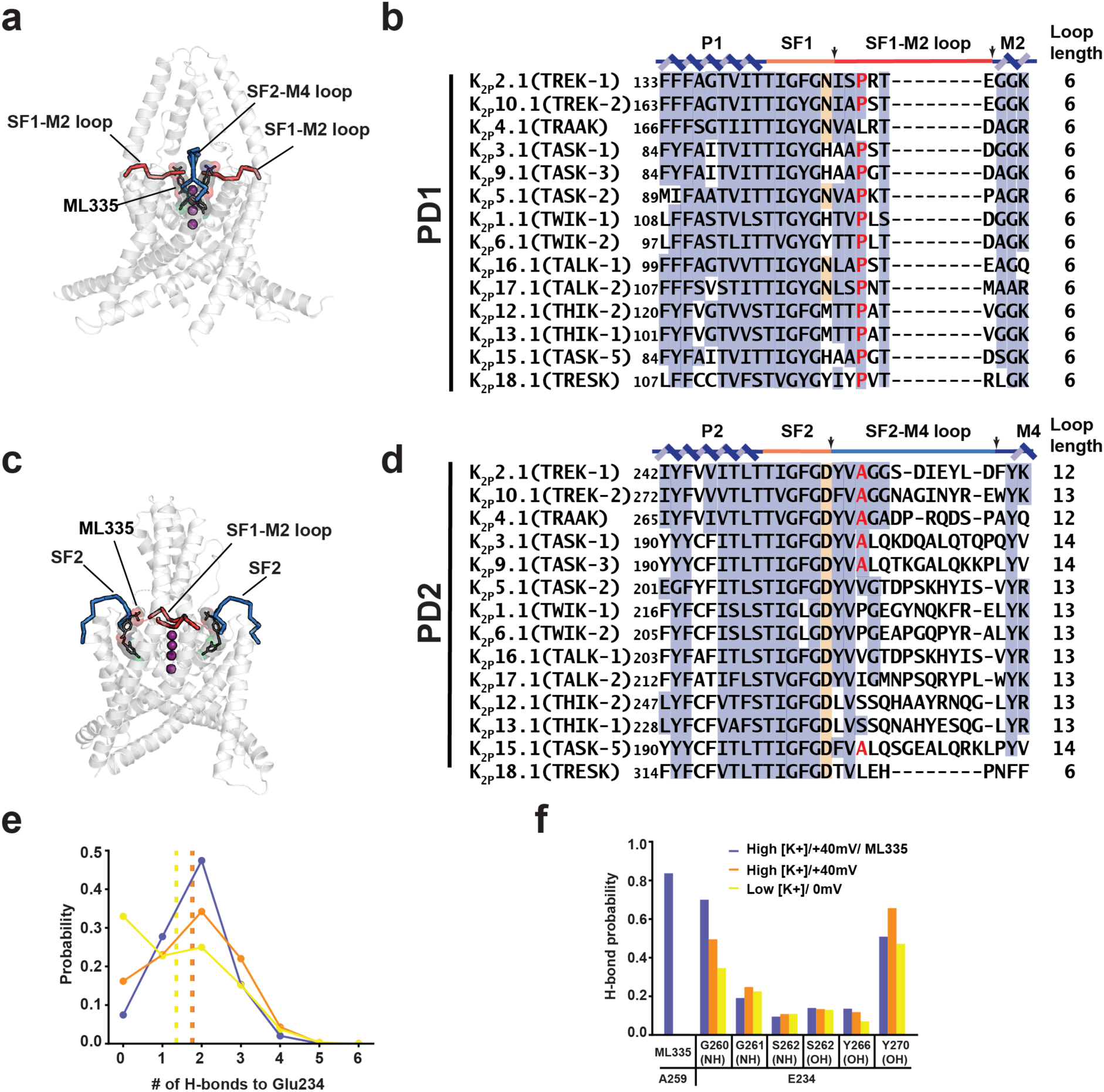
K_2P_ channel pore domain comparisons. **a**, K_2P_2.1 (TREK-1):ML335 complex (white) with a view showing the SF1-M2 loop. SF1-M2 loop (red) and SF2-M4 loop (blue) are indicted. ML335 (black) is shown in sticks with a transparent surface. **b**, Sequence alignment of PD1 for the indicated channels. P1 and M2 helices (blue), SF1 (orange), and SF1-M2 loop (red) are indicated. Terminal residue of the selectivity filter is highlighted. Arrows denote the boundaries of the SF1-M2 loop. **c**, K_2P_2.1 (TREK-1):ML335 complex (white) with a view showing the SF2-M4 loop. SF1-M2 loop (red) and SF2-M4 loop (blue) are indicted. ML335 (black) is shown in sticks with a transparent surface. **d**, Sequence alignment of PD2 for the indicated channels. Conserved residues are shaded in slate. Dashed red and blue boxes indicate the SF1-M2 (**c**) and SF2-M4 loops (**d**). Conserved selectivity filter N/D is shaded orange. Pro150, Ala259, and equivalents are shaded red. **e**, Per-frame probability of finding a particular number of hydrogen bonds to the Glu234 sidechain carboxylate in all K_2P_2.1 (TREK-1) simulations. Dotted lines indicate the overall average number of hydrogen bonds calculated for each simulation condition. **f**, Per-frame probability of a hydrogen bond between the indicated groups in all K_2P_2.1 (TREK-1) simulations. Sequences (**b,d**) are from human K_2P_ channels: K_2P_2.1 (TREK-1) AAD47569.1, K_2P_10.1 (TREK-2) NP_612190.1, K_2P_4.1 (TRAAK) AAI10328.1, K_2P_3.1 (TASK-1) NP_002237.1, K_2P_9.1 (TASK-3) NP_001269463.1, K_2P_5.1 (TASK-2) NP_003731.1, K_2P_1.1(TWIK-1) NP_002236.1, K_2P_6.1 (TWIK-2) NP_004814.1, K_2P_16.1 (TALK-1) NP_115491.1, K_2P_17.1 (TALK-2) AAK28551.1, K_2P_12.1 (THIK-2) NP_071338.1, K_2P_13.1 (THIK-1) NP_071337.2, K_2P_15.1 (TASK-5) EAW75900.1, K_2P_18.1 (TRESK) NP_862823.1. SF1 and SF2 sequence and numbers for K_2P_2.1 (TREK-1)_cryst_ (PDB:6CQ6)^7^ are identical to that of K_2P_2.1 (TREK-1) AAD47569.1.

**Fig. S7.**
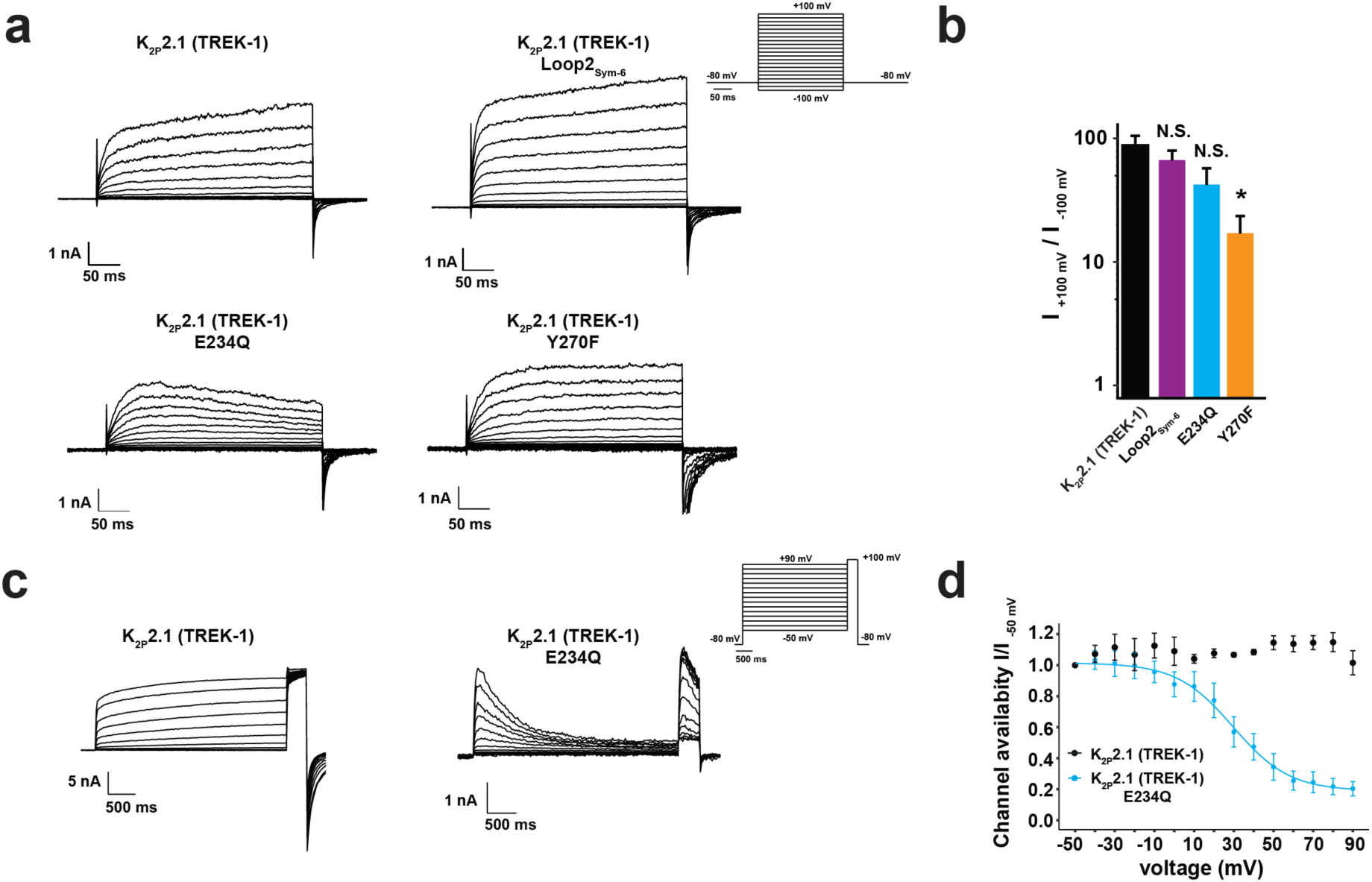
K_2P_ patch clamp recordings. **a**, Exemplar current traces from inside-out membrane patches of HEK293 cells expressing K_2P_2.1 (TREK-1), K_2P_2.1 (TREK-1) E234Q, K_2P_2.1 (TREK-1) Y270F, or K_2P_2.1 (TREK1) Loop2_Sym6_, in 150 mM K^+^_[ext]_/150 mM Rb^+^_[int]_. Inset shows voltage protocol. **b**, Rectification coefficients (I_+100mV_/-I_-100mV_) calculated from currents recorded on n ≥ 5 individual patches. * p < 0.05 compared to K_2P_2.1 (TREK-1). N.S, not statistically different. **c**, Exemplar current traces from inside-out membrane patches of HEK293 cells expressing K_2P_2.1 (TREK-1) or K_2P_2.1 (TREK-1) E234Q in response to an inactivation protocol (inset). **d**, Channel availability curves determined by plotting the normalized peak currents (I/I_-50mV_) measured at +100 mV as a function of pre-pulse voltages (n≥4). For panels **b**, and **d**, data represent mean ± s.e.m.

**Fig. S8.**
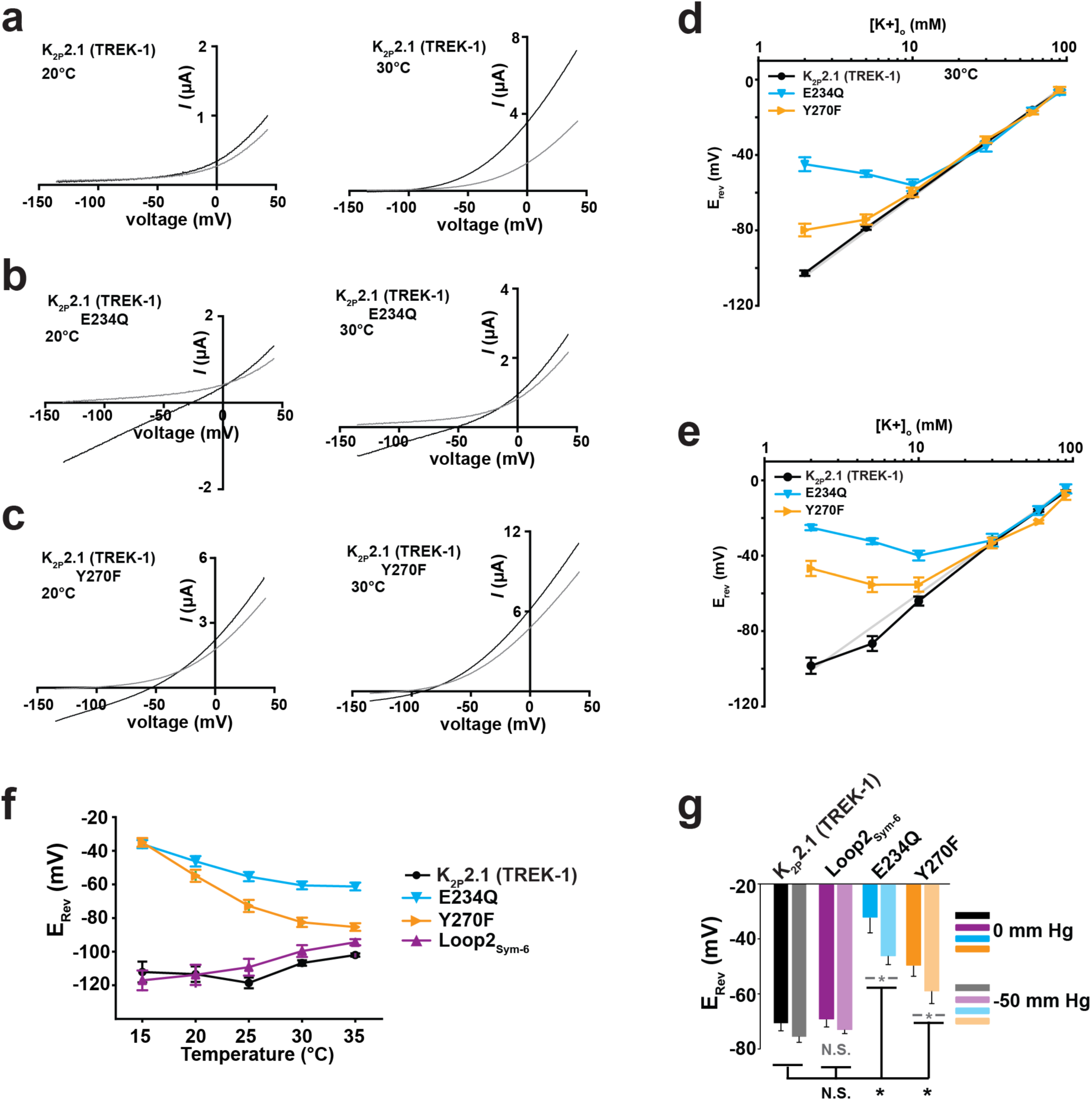
Activation alters K_2P_2.1 (TREK-1) mutant ion selectivity. **a-c**, Exemplar two-electrode voltage clamp (TEVC) recordings of K_2P_2.1 (TREK-1) (**a**), K_2P_2.1 (TREK-1) E234Q (**b**), and K_2P_2.1 (TREK-1) Y270F (**c**) at 20°C (left) and 30°C (right) in solutions of 96 mM Na^+^/2 mM K^+^ (black) or 96 mM *N*-methyl-D-glucamine/2 mM K^+^ (grey). **d,e**, Potassium selectivity recorded in *Xenopus* oocytes in K^+^/Na^*+*^ solutions (98.0 mM total) at pH_o_ = 7.4 at 20°C (**d**) and 30°C (**e**) (n=6). Data are background subtracted using uninjected oocytes. Grey line represents Nernst equation *E*_rev_ = *RT*/*F* × log([K^+^]_o_/[K^+^]_i_), where *R* and *F* have their usual thermodynamic meanings, *z* is equal to 1, and *T* = 20°C or 30°C, assuming [K^+^]_i_ = 108.6 mM (^89^). **f**, E_rev_ as a function of temperature for the indicated channels from TEVC experiments in *Xenopus* oocytes. **g**, E_rev_ at 0 and -50 mmHg measured from inside-out membrane patches of HEK293 cells expressing the indicated channels. * p < 0.05 compared to K_2P_2.1 (TREK-1) at the same pressure. N.S, not statistically different. (n ≥ 4). Grey indicates statistical significance between the 0 mM and -50 mM Hg measurements. For panels (**d**)-(**g**), data represent mean ± s.e.m.

**Supplementary Video 1 High [K**^**+**^**]/+40 mV simulation trajectory**. Pore helix 1 and SF1-M2 loop are shown in cartoon representation, SF1 is shown as sticks, and potassium ions are spheres of varying colors. Video represents the first 1920 ns of simulation 1.

**Supplementary Video 2 Low [K**^**+**^**]/0 mV simulation trajectory**. Pore helix 2 and SF2-M4 loop are shown in cartoon representation, SF2 is shown as sticks, M4 is shown as transparent cartoon, and the potassium ion is shown as a magenta sphere. Residues involved in the Glu234 network are shown as green sticks. Video represents the first 1920 ns of simulation 29.

**Supplementary Video 3 Morph between the SF1 active and inactive conformations**. 0 mM [K^+^]:ML335 structures and 1 mM [K^+^] structures represent the active and inactive conformations, respectively. Selectivity filter is yellow orange. Asn147 and Thr142 are shown as sticks. Potassium ions are magenta spheres.

**Supplementary Video 4 Morph between the SF2 active and inactive conformations**. 0 mM [K^+^]:ML335 structures and 1 mM [K^+^] structures represent the active and inactive conformations, respectively. Selectivity filter is yellow orange. Asp256 and Thr251 are shown as sticks. Potassium ions are magenta spheres.

**Table S1.**
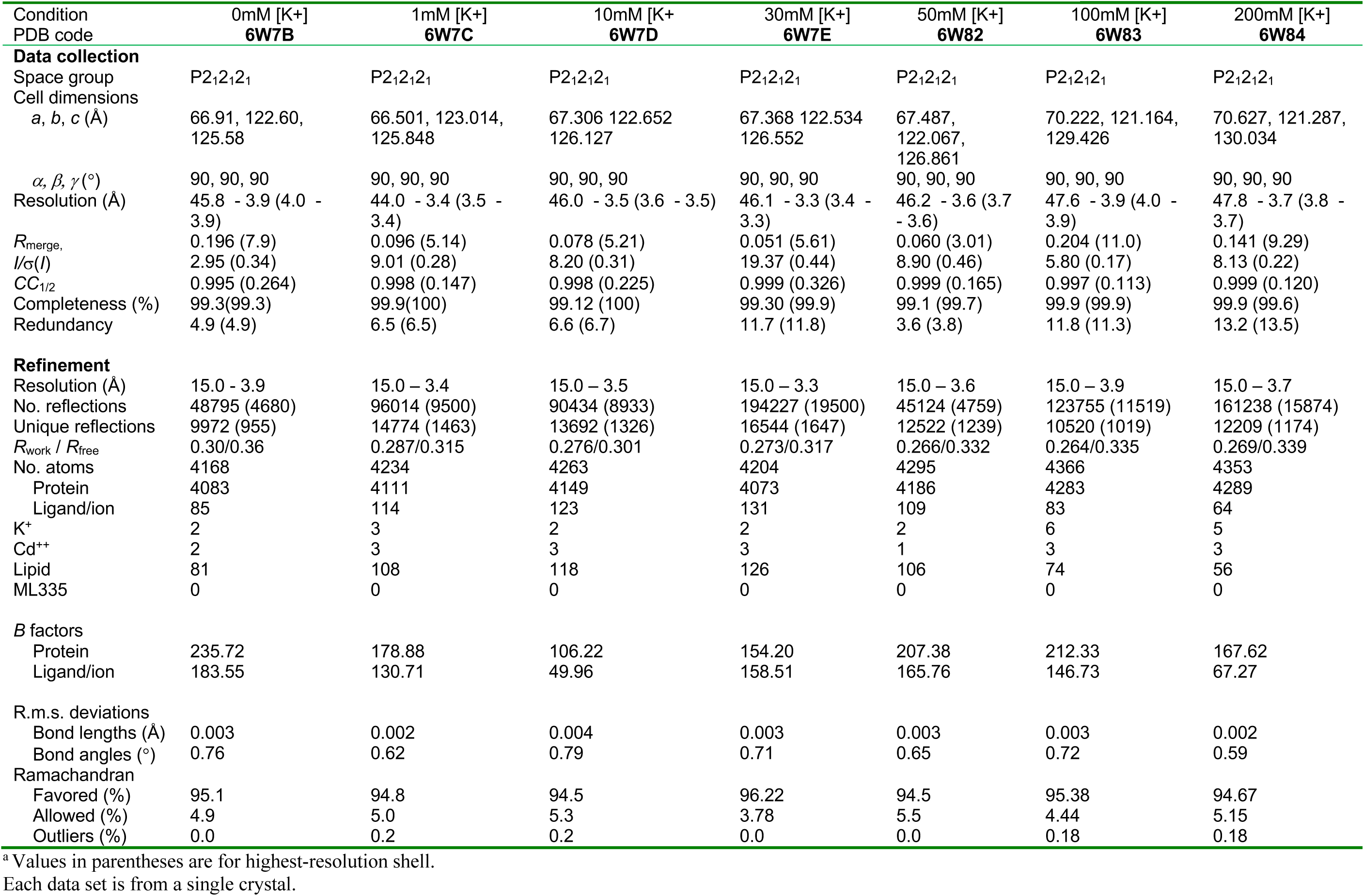
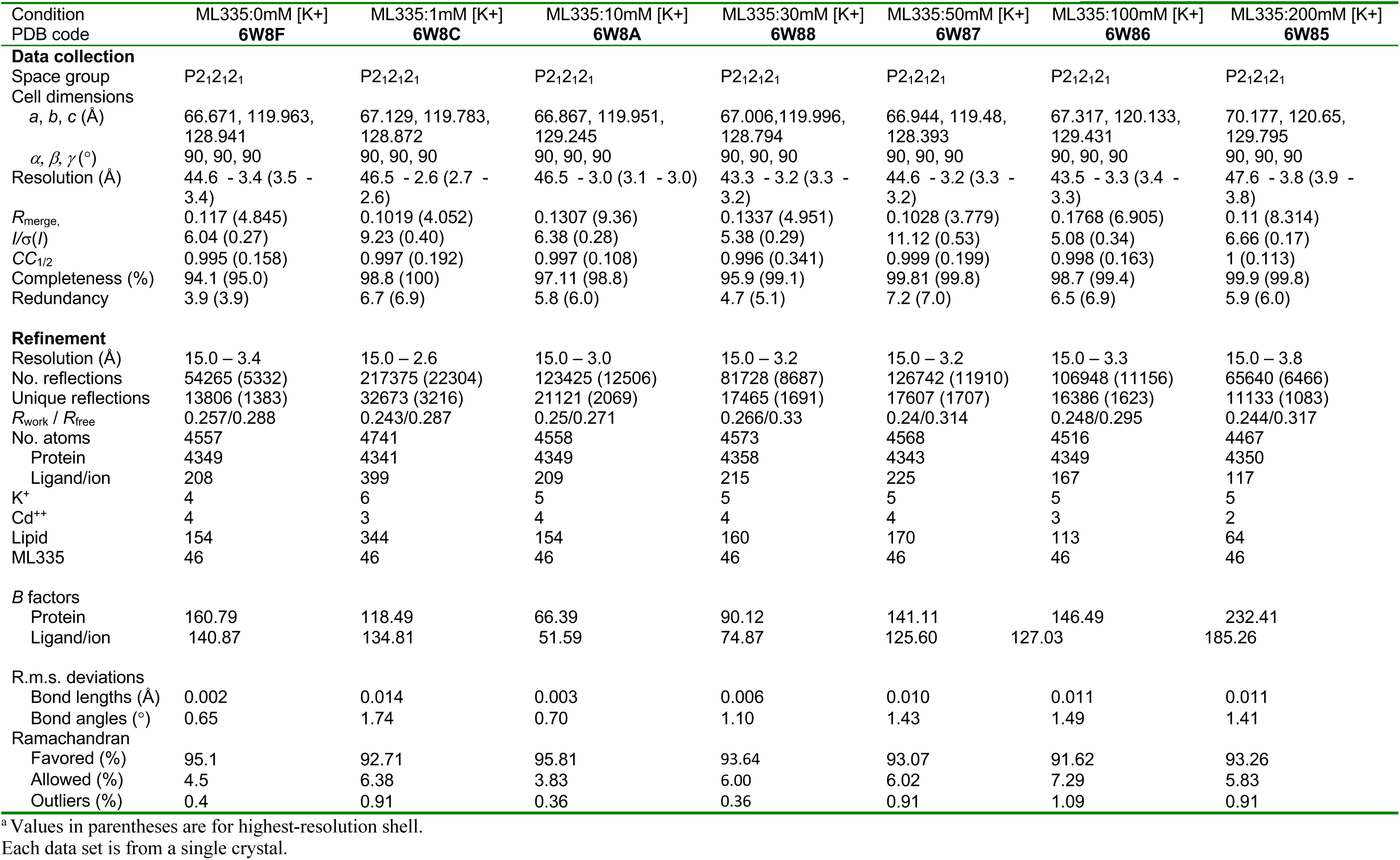
Data collection and refinement statistics K_2P_2.1 (TREK-1) and K_2P_2.1 (TREK-1):ML335 complex structures

| Condition<br>PDB code | 0mM [K+]<br>**6W7B** | 1mM [K+]<br>**6W7C** | 10mM [K+]<br>**6W7D** | 30mM [K+]<br>**6W7E** | 50mM [K+]<br>**6W82** | 100mM [K+]<br>**6W83** | 200mM [K+]<br>**6W84** |
| --- | --- | --- | --- | --- | --- | --- | --- |
| **Data collection** |  |  |  |  |  |  |  |
| Space group | P2<sub>1</sub>2<sub>1</sub>2<sub>1</sub> | P2<sub>1</sub>2<sub>1</sub>2<sub>1</sub> | P2<sub>1</sub>2<sub>1</sub>2<sub>1</sub> | P2<sub>1</sub>2<sub>1</sub>2<sub>1</sub> | P2<sub>1</sub>2<sub>1</sub>2<sub>1</sub> | P2<sub>1</sub>2<sub>1</sub>2<sub>1</sub> | P2<sub>1</sub>2<sub>1</sub>2<sub>1</sub> |
| Cell dimensions<br>a, b, c (Å) | 66.91, 122.60,<br>125.58 | 66.501, 123.014,<br>125.848 | 67.306 122.652<br>126.127 | 67.368 122.534<br>126.552 | 67.487,<br>122.067,<br>126.861 | 70.222, 121.164,<br>129.426 | 70.627, 121.287,<br>130.034 |
| $\alpha, \beta, \gamma$ (°) | 90, 90, 90 | 90, 90, 90 | 90, 90, 90 | 90, 90, 90 | 90, 90, 90 | 90, 90, 90 | 90, 90, 90 |
| Resolution (Å) | 45.8 - 3.9 (4.0 -<br>3.9) | 44.0 - 3.4 (3.5 -<br>3.4) | 46.0 - 3.5 (3.6 - 3.5) | 46.1 - 3.3 (3.4 -<br>3.3) | 46.2 - 3.6 (3.7<br>- 3.6) | 47.6 - 3.9 (4.0 -<br>3.9) | 47.8 - 3.7 (3.8 -<br>3.7) |
| *R*<sub>merge</sub>, | 0.196 (7.9) | 0.096 (5.14) | 0.078 (5.21) | 0.051 (5.61) | 0.060 (3.01) | 0.204 (11.0) | 0.141 (9.29) |
| *I*/$\sigma$(*I*) | 2.95 (0.34) | 9.01 (0.28) | 8.20 (0.31) | 19.37 (0.44) | 8.90 (0.46) | 5.80 (0.17) | 8.13 (0.22) |
| *CC*<sub>1/2</sub> | 0.995 (0.264) | 0.998 (0.147) | 0.998 (0.225) | 0.999 (0.326) | 0.999 (0.165) | 0.997 (0.113) | 0.999 (0.120) |
| Completeness (%) | 99.3(99.3) | 99.9(100) | 99.12 (100) | 99.30 (99.9) | 99.1 (99.7) | 99.9 (99.9) | 99.9 (99.6) |
| Redundancy | 4.9 (4.9) | 6.5 (6.5) | 6.6 (6.7) | 11.7 (11.8) | 3.6 (3.8) | 11.8 (11.3) | 13.2 (13.5) |
| **Refinement** |  |  |  |  |  |  |  |
| Resolution (Å) | 15.0 - 3.9 | 15.0 - 3.4 | 15.0 - 3.5 | 15.0 - 3.3 | 15.0 - 3.6 | 15.0 - 3.9 | 15.0 - 3.7 |
| No. reflections | 48795 (4680) | 96014 (9500) | 90434 (8933) | 194227 (19500) | 45124 (4759) | 123755 (11519) | 161238 (15874) |
| Unique reflections | 9972 (955) | 14774 (1463) | 13692 (1326) | 16544 (1647) | 12522 (1239) | 10520 (1019) | 12209 (1174) |
| *R*<sub>work</sub> / *R*<sub>free</sub> | 0.30/0.36 | 0.287/0.315 | 0.276/0.301 | 0.273/0.317 | 0.266/0.332 | 0.264/0.335 | 0.269/0.339 |
| No. atoms | 4168 | 4234 | 4263 | 4204 | 4295 | 4366 | 4353 |
| Protein | 4083 | 4111 | 4149 | 4073 | 4186 | 4283 | 4289 |
| Ligand/ion | 85 | 114 | 123 | 131 | 109 | 83 | 64 |
| K<sup>+</sup> | 2 | 3 | 2 | 2 | 2 | 6 | 5 |
| Cd<sup>++</sup> | 2 | 3 | 3 | 3 | 1 | 3 | 3 |
| Lipid | 81 | 108 | 118 | 126 | 106 | 74 | 56 |
| ML335 | 0 | 0 | 0 | 0 | 0 | 0 | 0 |
| **B factors** |  |  |  |  |  |  |  |
| Protein | 235.72 | 178.88 | 106.22 | 154.20 | 207.38 | 212.33 | 167.62 |
| Ligand/ion | 183.55 | 130.71 | 49.96 | 158.51 | 165.76 | 146.73 | 67.27 |
| **R.m.s. deviations** |  |  |  |  |  |  |  |
| Bond lengths (Å) | 0.003 | 0.002 | 0.004 | 0.003 | 0.003 | 0.003 | 0.002 |
| Bond angles (°) | 0.76 | 0.62 | 0.79 | 0.71 | 0.65 | 0.72 | 0.59 |
| **Ramachandran** |  |  |  |  |  |  |  |
| Favored (%) | 95.1 | 94.8 | 94.5 | 96.22 | 94.5 | 95.38 | 94.67 |
| Allowed (%) | 4.9 | 5.0 | 5.3 | 3.78 | 5.5 | 4.44 | 5.15 |
| Outliers (%) | 0.0 | 0.2 | 0.2 | 0.0 | 0.0 | 0.18 | 0.18 |

<sup>a</sup> Values in parentheses are for highest-resolution shell.

| Condition<br>PDB code | ML335:0mM [K+]<br>**6W8F** | ML335:1mM [K+]<br>**6W8C** | ML335:10mM [K+]<br>**6W8A** | ML335:30mM [K+]<br>**6W88** | ML335:50mM [K+]<br>**6W87** | ML335:100mM [K+]<br>**6W86** | ML335:200mM [K+]<br>**6W85** |
| --- | --- | --- | --- | --- | --- | --- | --- |
| **Data collection** |  |  |  |  |  |  |  |
| Space group | P2<sub>1</sub>2<sub>1</sub>2<sub>1</sub> | P2<sub>1</sub>2<sub>1</sub>2<sub>1</sub> | P2<sub>1</sub>2<sub>1</sub>2<sub>1</sub> | P2<sub>1</sub>2<sub>1</sub>2<sub>1</sub> | P2<sub>1</sub>2<sub>1</sub>2<sub>1</sub> | P2<sub>1</sub>2<sub>1</sub>2<sub>1</sub> | P2<sub>1</sub>2<sub>1</sub>2<sub>1</sub> |
| Cell dimensions<br>a, b, c (Å) | 66.671, 119.963,<br>128.941 | 67.129, 119.783,<br>128.872 | 66.867, 119.951,<br>129.245 | 67.006, 119.996,<br>128.794 | 66.944, 119.48,<br>128.393 | 67.317, 120.133,<br>129.431 | 70.177, 120.65,<br>129.795 |
| $\alpha, \beta, \gamma$ (°) | 90, 90, 90 | 90, 90, 90 | 90, 90, 90 | 90, 90, 90 | 90, 90, 90 | 90, 90, 90 | 90, 90, 90 |
| Resolution (Å) | 44.6 - 3.4 (3.5 -<br>3.4) | 46.5 - 2.6 (2.7 -<br>2.6) | 46.5 - 3.0 (3.1 - 3.0) | 43.3 - 3.2 (3.3 -<br>3.2) | 44.6 - 3.2 (3.3 -<br>3.2) | 43.5 - 3.3 (3.4 -<br>3.3) | 47.6 - 3.8 (3.9 -<br>3.8) |
| $R_{\text{merge}}$ | 0.117 (4.845) | 0.1019 (4.052) | 0.1307 (9.36) | 0.1337 (4.951) | 0.1028 (3.779) | 0.1768 (6.905) | 0.11 (8.314) |
| $I/\sigma(I)$ | 6.04 (0.27) | 9.23 (0.40) | 6.38 (0.28) | 5.38 (0.29) | 11.12 (0.53) | 5.08 (0.34) | 6.66 (0.17) |
| $CC_{1/2}$ | 0.995 (0.158) | 0.997 (0.192) | 0.997 (0.108) | 0.996 (0.341) | 0.999 (0.199) | 0.998 (0.163) | 1 (0.113) |
| Completeness (%) | 94.1 (95.0) | 98.8 (100) | 97.11 (98.8) | 95.9 (99.1) | 99.81 (99.8) | 98.7 (99.4) | 99.9 (99.8) |
| Redundancy | 3.9 (3.9) | 6.7 (6.9) | 5.8 (6.0) | 4.7 (5.1) | 7.2 (7.0) | 6.5 (6.9) | 5.9 (6.0) |
| **Refinement** |  |  |  |  |  |  |  |
| Resolution (Å) | 15.0 - 3.4 | 15.0 - 2.6 | 15.0 - 3.0 | 15.0 - 3.2 | 15.0 - 3.2 | 15.0 - 3.3 | 15.0 - 3.8 |
| No. reflections | 54265 (5332) | 217375 (22304) | 123425 (12506) | 81728 (8687) | 126742 (11910) | 106948 (11156) | 65640 (6466) |
| Unique reflections | 13806 (1383) | 32673 (3216) | 21121 (2069) | 17465 (1691) | 17607 (1707) | 16386 (1623) | 11133 (1083) |
| $R_{\text{work}} / R_{\text{free}}$ | 0.257/0.288 | 0.243/0.287 | 0.25/0.271 | 0.266/0.33 | 0.24/0.314 | 0.248/0.295 | 0.244/0.317 |
| No. atoms | 4557 | 4741 | 4558 | 4573 | 4568 | 4516 | 4467 |
| Protein | 4349 | 4341 | 4349 | 4358 | 4343 | 4349 | 4350 |
| Ligand/ion | 208 | 399 | 209 | 215 | 225 | 167 | 117 |
| K<sup>+</sup> | 4 | 6 | 5 | 5 | 5 | 5 | 5 |
| Cd<sup>++</sup> | 4 | 3 | 4 | 4 | 4 | 3 | 2 |
| Lipid | 154 | 344 | 154 | 160 | 170 | 113 | 64 |
| ML335 | 46 | 46 | 46 | 46 | 46 | 46 | 46 |
| **B factors** |  |  |  |  |  |  |  |
| Protein | 160.79 | 118.49 | 66.39 | 90.12 | 141.11 | 146.49 | 232.41 |
| Ligand/ion | 140.87 | 134.81 | 51.59 | 74.87 | 125.60 | 127.03 | 185.26 |
| **R.m.s. deviations** |  |  |  |  |  |  |  |
| Bond lengths (Å) | 0.002 | 0.014 | 0.003 | 0.006 | 0.010 | 0.011 | 0.011 |
| Bond angles (°) | 0.65 | 1.74 | 0.70 | 1.10 | 1.43 | 1.49 | 1.41 |
| **Ramachandran** |  |  |  |  |  |  |  |
| Favored (%) | 95.1 | 92.71 | 95.81 | 93.64 | 93.07 | 91.62 | 93.26 |
| Allowed (%) | 4.5 | 6.38 | 3.83 | 6.00 | 6.02 | 7.29 | 5.83 |
| Outliers (%) | 0.4 | 0.91 | 0.36 | 0.36 | 0.91 | 1.09 | 0.91 |

<sup>a</sup> Values in parentheses are for highest-resolution shell.
Each data set is from a single crystal.

**Table S2.** Anomalous peak heights (in σ)

|  | 1 mM [K<sup>+</sup>] | 200mM [K<sup>+</sup>] | ML335:1mM [K<sup>+</sup>] | ML335:200mM [K<sup>+</sup>] |
| --- | --- | --- | --- | --- |
| **Filter site** |  |  |  |  |
| S1 | 2.25 | 6.24 | 5.47 | 6.09 |
| S2 | 3.10 | 7.64 | 6.12 | 7.01 |
| S3 | 6.24 | 8.30 | 6.40 | 8.23 |
| S4 | 3.89 | 6.19 | 4.38 | 4.09 |
| Resolution (Å) | 5.5 | 5.7 | 5.0 | 5.5 |

Anomalous peak heights (in $\sigma$ ) from long-wavelength data above ( $\lambda = 3.35$ Å) the potassium K-edge as calculated with ANODE<sup>65</sup> based on K<sub>2</sub>P2.1 (TREK-1) (6CQ6)<sup>7</sup>.

**Table S3.** Molecular dynamics simulations.

| ID | PDBID | n atoms | Engine | [K<sup>+</sup>] (mM) | ML335 | Potential (mV) | Length (ns) | n permeations |
| --- | --- | --- | --- | --- | --- | --- | --- | --- |
| 1 | 6W8C | 205060 | Anton2 | 180 | yes | +40 | 2880 | 26 |
| 2 | 6W8C | 205060 | Anton2 | 180 | yes | +40 | 2880 | 23 |
| 3 | 6W8C | 205060 | Anton2 | 180 | yes | +40 | 2880 | 9 |
| 4 | 6W8C | 205060 | Anton2 | 180 | yes | +40 | 3840 | 30 |
| 5 | 6W8C | 205060 | Anton2 | 180 | yes | +40 | 2880 | 35 |
| 6 | 6W8C | 205228 | Anton2 | 180 | yes | +40 | 3840 | 26 |
| 7 | 6W8C | 205228 | Anton2 | 180 | yes | +40 | 2880 | 20 |
| 8 | 6W8C | 205228 | Anton2 | 180 | yes | +40 | 2880 | 2 |
| 9 | 6W8C | 205228 | Anton2 | 180 | yes | +40 | 2880 | 29 |
| 10 | 6W8C | 205228 | Anton2 | 180 | yes | +40 | 3840 | 53 |
| 11 | 5VK5 | 205159 | Anton2 | 180 | no | +40 | 4800 | 3 |
| 12 | 5VK5 | 205159 | Anton2 | 180 | no | +40 | 4800 | 36 |
| 13 | 6CQ6 | 205175 | Anton2 | 180 | no | +40 | 1920 | 3 |
| 14 | 6CQ6 | 205175 | Anton2 | 180 | no | +40 | 1920 | 2 |
| 15 | 6CQ6 | 205175 | Anton2 | 180 | no | +40 | 1920 | 2 |
| 16 | 6CQ6 | 205175 | Anton2 | 180 | no | +40 | 1920 | -1 |
| 17 | 6CQ6 | 205175 | Anton2 | 180 | no | +40 | 2880 | 19 |
| 18 | 6CQ6 | 205325 | Anton2 | 180 | no | +40 | 3840 | 20 |
| 19 | 6CQ6 | 205325 | Anton2 | 180 | no | +40 | 2880 | 18 |
| 20 | 6CQ6 | 205325 | Anton2 | 180 | no | +40 | 2880 | 6 |
| 21 | 6CQ6 | 205325 | Anton2 | 180 | no | +40 | 2880 | 2 |
| 22 | 6CQ6 | 205325 | Anton2 | 180 | no | +40 | 3840 | 34 |
| 23 | 6CQ6 | 204174 | gromacs | 5 | no | 0 | 1400 | - |
| 24 | 6CQ6 | 204168 | gromacs | 5 | no | 0 | 2400 | - |
| 25 | 6CQ6 | 204168 | gromacs | 5 | no | 0 | 2000 | - |
| 26 | 6CQ6 | 204006 | gromacs | 5 | no | 0 | 2000 | - |
| 27 | 6CQ6 | 204006 | gromacs | 5 | no | 0 | 2000 | - |
| 28 | 6CQ6 | 204174 | Anton2 | 5 | no | 0 | 3600 | - |
| 29 | 6CQ6 | 204441 | Anton2 | 5 | no | 0 | 3600 | - |
| 30 | 6CQ6 | 204006 | Anton2 | 5 | no | 0 | 3600 | - |

